# Non-catalytic regulation of 18*S* rRNA methyltransferase DIMT1 in acute myeloid leukemia

**DOI:** 10.1101/2023.03.18.533235

**Authors:** Yulia Gonskikh, Julian Stoute, Hui Shen, Krista Budinich, Bianca Pingul, Kollin Schultz, Heidi Elashal, Ronen Marmorstein, Junwei Shi, Kathy Fange Liu

## Abstract

Several rRNA modifying enzymes install rRNA modifications while participating in ribosome assembly. Here, we show that 18*S* rRNA methyltransferase DIMT1 is essential for acute myeloid leukemia (AML) proliferation through a non-catalytic function. We reveal that targeting a positively charged cleft of DIMT1, remote from the catalytic site, weakens the binding of DIMT1 to rRNA and mis-localizes DIMT1 to the nucleoplasm, in contrast with the primarily nucleolar localization of wild-type DIMT1. Mechanistically, rRNA binding is required for DIMT1 to undergo liquid-liquid phase separation, which explains why the distinct nucleoplasm localization of the rRNA-binding deficient DIMT1. Re-expression of wild-type or a catalytically inactive mutant E85A, but not the rRNA-binding deficient DIMT1, supports AML cell proliferation. This study provides a new strategy to target DIMT1-regulated AML proliferation via targeting this essential non-catalytic region.

## Introduction

Dimethyladenosine transferase 1 (DIMT1), a *S*-adenosyl methionine (SAM)-dependent methyltransferase, is an rRNA modifying enzyme which participates in ribosome biogenesis (Chaker-Margot et al. 2015; Chaker-Margot et al. 2017; Klinge and Woolford 2019). DIMT1 installs *N*^6,6^-dimethyladenosine (m_2_^6,6^A) at the two adjacent adenosine sites A1850 and A1851, almost to 100% occupancy, in human 18*S* rRNA (Poldermans et al. 1980; O’Farrell et al. 2004; Mangat and Brown 2008; Tu et al. 2009; Boehringer et al. 2012). Recent studies reveal that the catalytic role of DIMT1 is important for stress response (Helser et al. 1971; Lafontaine et al. 1994; Tokuhisa et al. 1998; Wieckowski and Schiefelbein 2012) and is required for translational fidelity in bacterial and human cells (Shen et al. 2020b). DIMT1 is located in the cell nucleolus where rRNA transcription, rRNA processing, and ribosome assembly take place (Zorbas et al. 2015). Specifically, DIMT1 participates in the late steps of 18*S* rRNA processing in a non-catalytical manner (Zorbas et al. 2015). Ablation of DIMT1 disrupts ribosome biogenesis, and it is lethal for human cells. Recently, we and others have shown that the catalytic activity of DIMT1 is not required for ribosome biogenesis (Zorbas et al. 2015; Shen et al. 2020b; Shen et al. 2021). Thus, the function of DIMT1-mediated rRNA methylation and DIMT1’s non-catalytic role in rRNA processing and ribosome biogenesis are decoupled.

Ribosome biogenesis is fundamentally important for cell growth and proliferation. Although ribosomes are ubiquitously expressed in all cell types, certain cell types (such as hematopoietic cells) are more affected by ribosomal defects than others (Mills and Green 2017). For instance, ribosomal defects are frequently seen in bone marrow failure, anemia, and hematopoietic malignancy. Mutations in ribosomal proteins RPL5 and RPL10 of the large ribosomal subunit have been implicated in T-cell acute lymphoblastic leukemia, and mutations in RPS15 of the small ribosomal subunit have been implicated in chronic lymphocytic leukemia (Vlachos 2017). Early ribosome biogenesis occurs in the cell nucleolus where represents a multilayered biomolecular condensate that is formed through a biophysical process called liquid-liquid phase separation (LLPS) (Feric et al. 2016). LLPS facilitates the initial steps of ribosome biogenesis and several other functions (Lafontaine et al. 2021). For instance, ribosomal proteins and rRNA modifying enzymes such as the box C/D small nucleolar RNA-associated methyltransferase fibrillarin (FBL) and nucleophosmin (NPM1) reside in different layers in the nucleolus to facilitate rRNA synthesis and ribosomal subunit maturation steps (Amin et al. 2008). However, it is not known whether DIMT1 undergoes LLPS to the nucleolus to facilitate ribosome maturation or rRNA methylation. Since DIMT1 aids ribosome assembly in the nucleolus, understanding the mechanism of how DIMT1 localizes to the nucleolus may provide a new strategy to regulate its role in ribosome assembly.

Additionally, a diverse set of chemical modifications are found within rRNAs (Desrosiers et al. 1974; Desrosiers et al. 1975; du Toit 2016; Gilbert et al. 2016; Lewis et al. 2017; Roundtree et al. 2017; Willyard 2017; Zhao et al. 2017). Critically, these modifications can dramatically alter rRNA structure, stability, and protein synthesis (Sloan et al. 2017). The functional significance of rRNA modifications is further evidenced by the fact that dysregulation of rRNA modifying enzymes and snoRNAs (which guide rRNA modifying enzymes to achieve substrate specificity) has been linked to a battery of human cancers, including hematopoietic malignancies (Narla and Ebert 2010; McMahon et al. 2015; Frye and Blanco 2016; Nachmani et al. 2019; Janin et al. 2020; Barros-Silva et al. 2021). DIMT1 is highly expressed in hematopoietic malignancies such as acute myeloid leukemia (AML) and multiple myeloma, and studies show that knockdown of DIMT1 leads to reduced tumorigenicity in myeloma (Zorbas et al. 2015; Ikeda et al. 2017; Ikeda et al. 2018; Janker et al. 2019). However, it is not known whether the catalytic role or the non-catalytic role of DIMT1 supports leukemia cell proliferation.

Here, we revealed a positively charged cleft that is remote from the catalytic center of DIMT1 and is essential for its RNA binding ability and function in ribosome biogenesis. Furthermore, this cleft is required for DIMT1 undergoing RNA-facilitated LLPS. Simultaneous mutation of those positively charged residues, which dampens the rRNA binding affinity of DIMT1, leads to aberrant nucleoplasm localization of DIMT1 and thus fails to support AML cell proliferation. The results also demonstrate the non-catalytic role of DIMT1 instead of DIMT1-mediated rRNA methylation is important for AML cell proliferation.

## Results

### DIMT1 depletion impairs 18*S* rRNA processing and AML cell proliferation

Previous studies have suggested that DIMT1 is highly expressed in leukemia cells (Shen et al. 2021) (https://depmap.org/portal/publications/). CERES score showed moderately higher dependency of leukemia cells on DIMT1 gene compare to all cancers (Supplementary Figure S1A). To validate the requirement of DIMT1 in cancer cell proliferation, we performed a competition-based proliferation assay across a selection of cancer cell lines. We transduced single-guide RNAs (sgRNAs) targeting DIMT1, Rosa (a negative control), and PCNA (a positive control) co-expressed with the GFP marker into two AML cell lines and two solid tumor cells lines (Supplementary Figure S1B). The resulting cell population was comprised of both transduced (GFP^+^) and untransduced cells (GFP^+^), and the relative abundance of each population was tracked via flow cytometry. Unlike the effects of sgRosa, we found that cells expressing sgDIMT1 were rapidly outcompeted by non-transduced cells, as shown by flow cytometry-based tracking of GFP expression. Furthermore, the cellular competition results showed that the AML cell lines (MOLM-13C and MV4-11C) preferentially required DIMT1 for proliferation in comparison to solid tumor cell lines (Figure 1A). The western blot analyses confirmed the successful depletion of DIMT1 after five days of sgDIMT1 administration (Figure 1B).

**Fig. 1.**
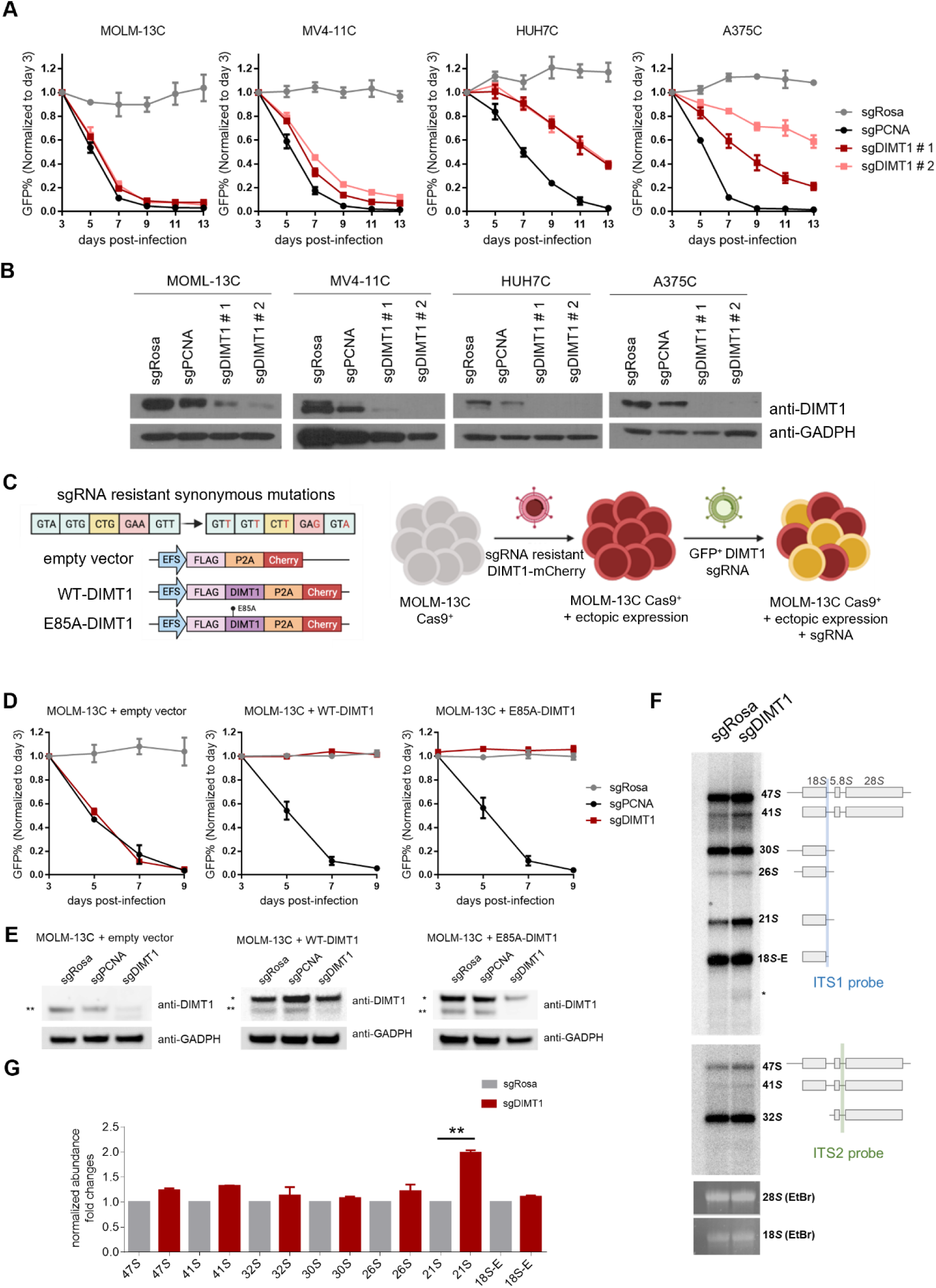
DIMT1 is essential for AML cell proliferation. (**A**) Competition-based proliferation assays performed in indicated Cas9^+^ cell lines. sgRNA^+^ populations were monitored over time with a GFP co-expression marker. Plotted is the relative sgRNA^+^ population normalized to the day 3 sgRNA^+^ population over 13 days. sgRosa is the negative control and sgPCNA is the positive control. (**B**) Western blots showing DIMT1 expression in cells transfected with the indicated sgRNAs on day 5 post-infection. (**C**) Schematic of re-expressing WT-DIMT1, E85A-DIMT1, or empty vector in MOLM-13C cells. DIMT1 carries synonymous mutations so it is not targeted by sgRNA of DIMT1. (**D**) Competition-based proliferation assay of MOLM-13C cells re-expressing WT-DIMT1, E85A-DIMT1, or an empty vector versus parental cells. (**E**) Western blots showing endogenous DIMT1 (annotated by two stars) and exogenous DIMT1 (annotated by one star) expression in cells transfected with the indicated sgRNAs on day 5 post-infection. (**F**) Northern blot analysis of rRNA precursors with total RNA extracted from MOLM13C cells transfected with negative control (sgRosa) or gRNA targeting DIMT1 (sgDIMT1). The detected pre-rRNA species are indicated to the right and schematized. The probes targeted ITS1 (upper panel) and ITS2 (lower panel). The mature 28*S* and 18*S* rRNAs were stained with ethidium bromide. (**G**) Quantification of rRNA precursors from Fig. 1F. The two-tailed t-test was used to calculate the p-value (p= 0.0012). Error bars represent mean ± s.d.

To further verify the essential function of DIMT1 and the on-target effect of our sgRNA, we ectopically expressed CRISPR-resistant WT-DIMT1 cDNA, which contains mutations at the sgRNA targeting site or an empty vector in leukemia cells (Figure 1C). To achieve a proper expression level of our transgenes, CRISPR-resistant WT-DIMT1, we transduced MOLM-13C cells at low multiplicity of infection (Figure S1C). Since exogenous DIMT1 had an mCherry tag separated by P2A linker, we performed fluorescence-activated cell sorting (FACS) to enrich cells expressing exogenous DIMT1. Low multiplicity of infection following by cell sorting allowed us to achieve expression of exogenous DIMT1 close to its endogenous level (expression levels of exogenous WT- and E85A DIMT1 are 2.69 ± 0.6 and 2.67 ± 0.4, while endogenous DIMT1 was normalized to 1). We then transfected the sgDIMT1 to target the endogenous DIMT1. As shown in Figure 1D, cells re-expressing WT-DIMT1 have rescued cell proliferation, but cells expressing the empty vector do not.

To dissect whether the enzymatic activity of DIMT1 is important for AML proliferation, we expressed CRISPR-resistant E85A-DIMT1 (catalytically inactive (Shen et al. 2020b)) in MOLM-13C cells and performed the same assays as WT-DIMT1. As shown in Figure 1 D and E, cells re-expressing E85A-DIMT1 rescued MOLM-13C cell proliferation. We further investigated whether DIMT1 depletion influences 18*S* rRNA processing by performing Northern blots. We utilized RNA probes targeting ITS1 (upper panel) and ITS2 (lower panel) to quantify the levels of the rRNA precursors. The results suggest that DIMT1 depletion caused a substantial accumulation of 21*S* pre-RNA compared to the control cells (Figure 1 F and G).

### DIMT1 depletion alters the expression and translation of transcripts involved in cell proliferation

To understand the impact of DIMT1 depletion on translation, we carried out ribosome profiling followed by high-throughput sequencing (ribo-seq) in MOLM-13C cells transfected with sgRosa versus cells transfected with sgDIMT1. The principal component analysis was summarized in Supplementary Figure S2A; the sequencing data of the ribosome protected RNA (RIBO) and the input RNA (RNA) from DIMT1 depletion, and the control cells separated well in sequencing group (RNA *vs.* RIBO) and treatment condition (sgRosa *vs.* sgDIMT1). The two biological replicates also cluster together, indicating the reliable quality of the sequencing data. Firstly, we analyzed the differential gene expression. We found that DIMT1 depletion leads to significantly increased expression of 134 genes (log_2_-fold change > 1) (Figure 2A, Supplementary Figure S2B, and Table S1); gene ontology (GO) analysis indicates the proteins encoded by these transcripts are mainly involved in the regulation of the immune response and cell adhesion (Figure 2B). DIMT1 depletion also significantly decreases the expression of 18 genes (log_2_-fold change < −1); gene ontology (GO) analysis indicates the proteins encoded by these transcripts are mainly involved in the regulation of the cell cycle (Figure 2A and B). Gene set enrichment analysis (GSEA) revealed that the MYC-target gene signature and the HOX gene cluster signature are significantly altered upon DIMT1 depletion indicating that dysregulation of DIMT1 may affect common cancer pathways and lead to a broad spectrum of cancers including leukemia (Figure 2C). Then, we analyzed the translational efficiency difference between DIMT1 depleted cells versus the control cells. As shown in Figure 2D, Supplementary Figure S2C, and Table S2, 70 transcripts exhibited significantly increased translation efficiency upon DIMT1 depletion; these transcripts are mainly involved in the regulation of leukocyte migration and spliceosome (Figure 2E). Another 46 transcripts showed decreased ribosome occupancies upon DIMT1 depletion; these transcripts are mainly involved in the regulation of translation and ribosome biogenesis, which is consistent with the function of DIMT1 as a ribosome assembly factor (Figure 2 D and E).

**Fig. 2.**
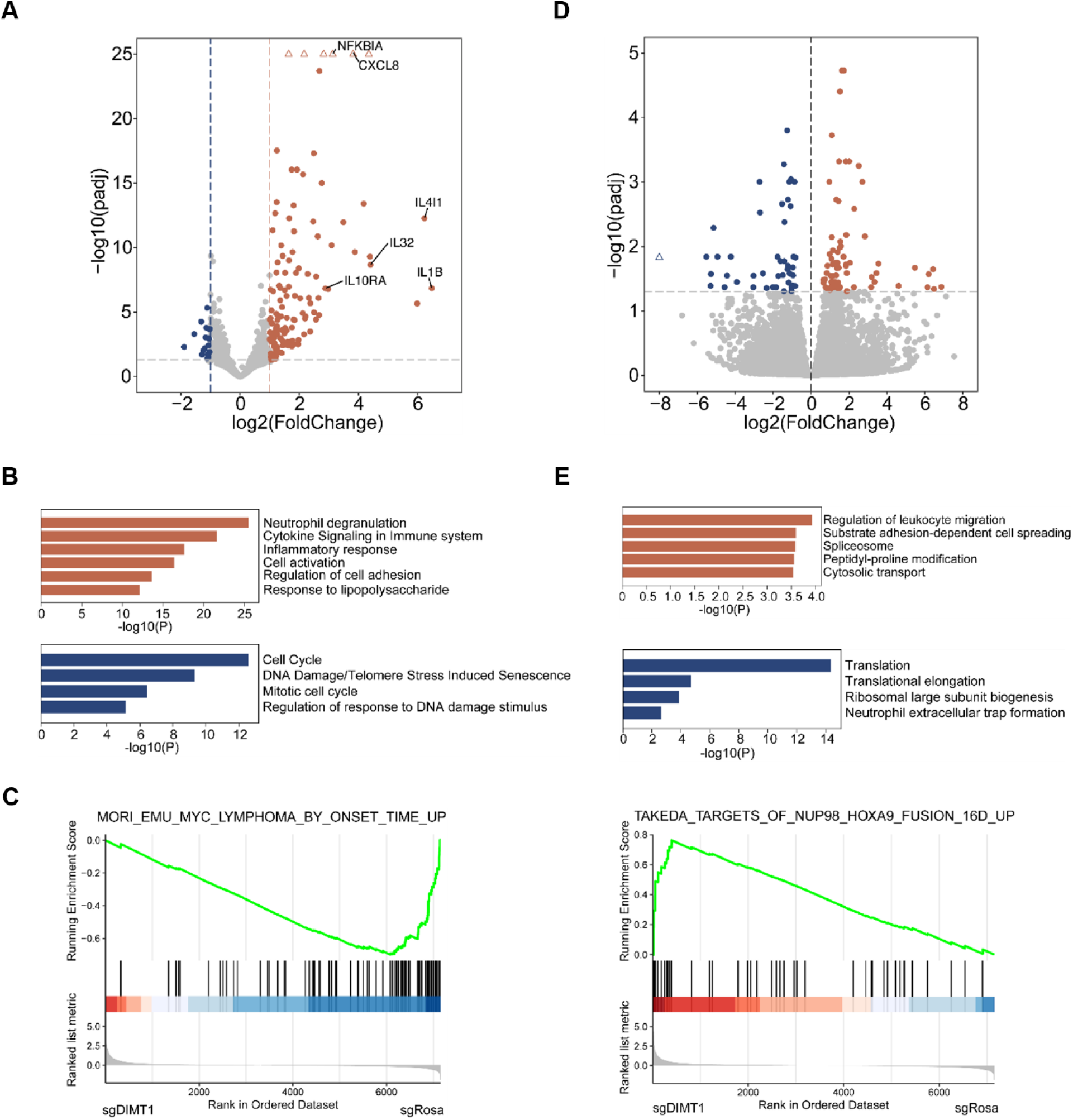
Effect of DIMT1 depletion on MOLM-13C cells. (**A**) A volcano plot shows the RNAs with differential expression levels in DIMT1 depleted versus control MOLM-13C cells. The significantly upregulated (padj < 0.05 and log_2_Foldchange > 1) transcripts are shown in orange color, while the significantly downregulated (padj < 0.05 and log_2_Foldchange < −1) transcripts are shown in blue color. (**B**) GO analysis of the significantly up and downregulated transcripts which are shown in Fig. 2A; transcripts with increased expression levels are in orange, and transcripts with decreased expression levels are in blue. (**C**) Gene set enrichment analysis (GSEA) of sequencing data presented in Fig. 2B. NUP98_HOXA9_fusion_up and MYC_lymphoma_up signatures were used. (**D**) A volcano plot shows the RNAs with differential translational efficiency in DIMT1 depleted versus control MOLM-13C cells. The significantly upregulated (padj < 0.05) genes are shown in orange color, while the significantly downregulated (padj < 0.05) genes are shown in blue color. (**E**) GO analysis of the differential transcripts in Fig. 6D; transcripts with increased translation efficiencies are in red, and transcripts with decreased translation efficiencies are in blue.

### A cleft in DIMT1 constituted of five positively charged residues is important for its RNA binding

DIMT1 associates with pre-ribosomes at the early steps of 18*S* rRNA processing and 40*S* small subunit assembly in the nucleolus (Zorbas et al. 2015). Since the catalytic activity of DIMT1 is not required for AML proliferation (Figure 1D), we next investigated whether we could impede DIMT1 joining ribosome assembly to reduce AML proliferation. We analyzed the residues in DIMT1 which contacts 18*S* rRNA. As shown in the structure of DIMT1 (PDB 7MQA) (Shen et al. 2021) (Figure 3 A and B), there is a positively charged cleft between the *N*-terminal domain and the *C*-terminal domain of DIMT1, which contacts helix 45 in 18*S* rRNA. There are five positively charged residues, including Arg162 and Arg174 in the *N*-terminal domain and Arg228, Lys253, and Arg256 in the C-terminal domain of DIMT1, which are particularly noteworthy for their potential rRNA binding ability (Figure 3B). We asked whether these residues contribute to DIMT1’s RNA binding and processing of 18*S* rRNA.

**Fig. 3.**
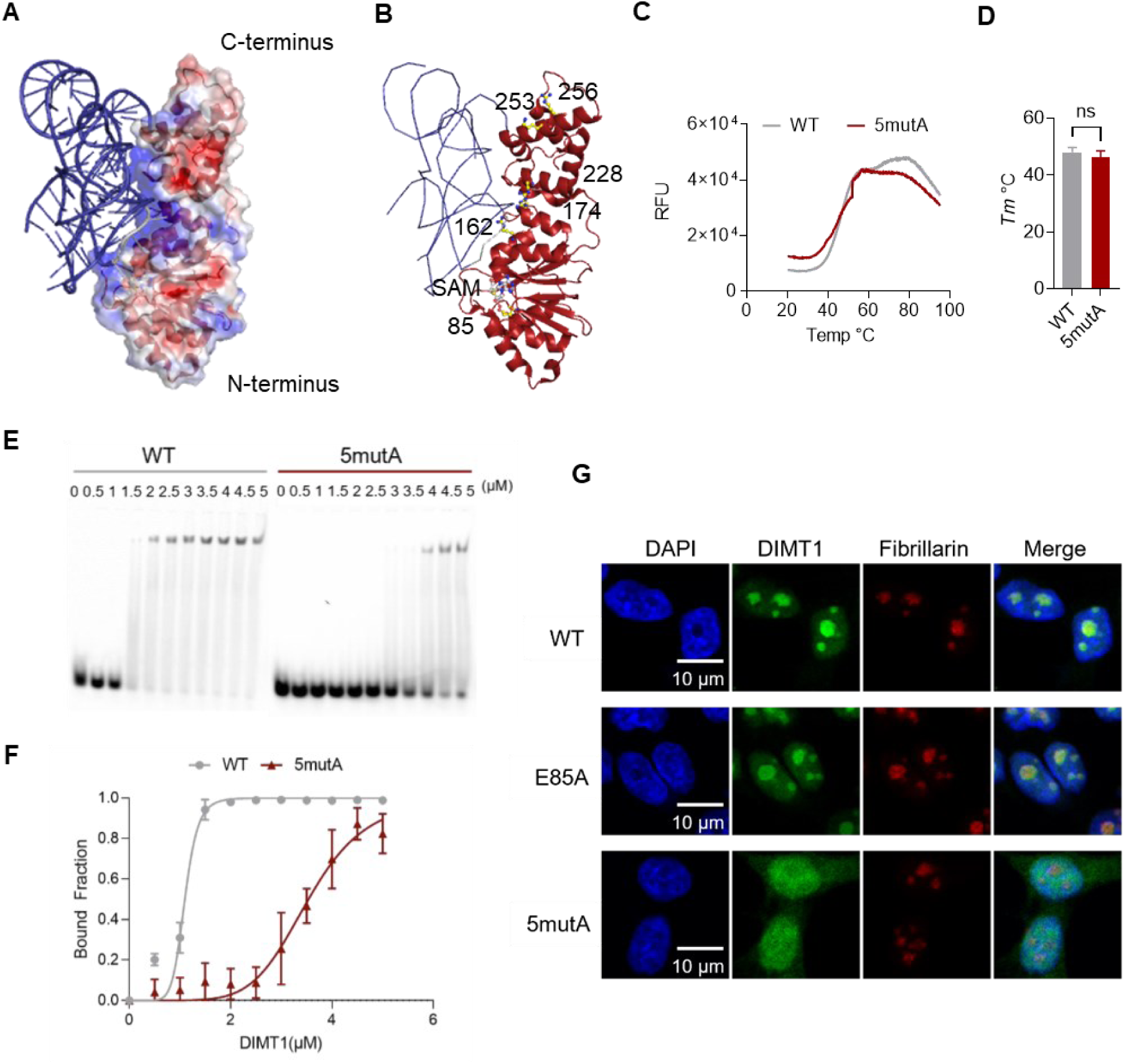
5mutA-DIMT1 presents weaker RNA-binding affinity. (**A**) The electrostatic surface of DIMT1 is shown to reveal the details of the RNA binding surface (PDB 7MQA). (**B**) Five positively charged residues, Glutamate 85, and SAM are depicted in stick and ball structures. (**C**) Differential scanning fluorometry (DSF) melting curves show the relative fluorescence units with increasing temperature. (**D**) The fluorescence is a product of SYPRO Orange dye, which resides on hydrophobic residues of exposed residues of WT- and 5mutA-DIMT1. The steady increase in fluorescence is due to the unfolding of the protein with increased temperature and the subsequent binding of the SYPRO Orange dye. (**E**) EMSA experiments show WT- and 5mutA-DIMT1 binding to the RNA probe. (**F**) RNA binding was plotted using the equation detailed in the Method section. WT-DIMT1 (Lanes 1 – 7, left) and 5mutA-DIMT1 (Lanes 8 – 14, right) at annotated concentrations and 5’ FAM labeled RNA probe at 1 µM were used. (**G**) Representative fluorescence microscopy images of Flag-tagged WT-, E85A-, and 5mutA-DIMT1-mClover in *DIMT1*^+/−^ HEK 293T cells. The nucleoli were stained with an anti-fibrillarin antibody as a marker (red), while the nuclei were stained with a NuclearMask dye (blue). Scale bar, 10 µm.

To investigate whether these positively charged residues in the cleft are important for DIMT1’s binding of 18*S* rRNA, we constructed a DIMT1 variant that contains five mutations inducing R162A, R174A, R228A, K253A, and R256A, which we refer to as 5mutA-DIMT1. We further expressed and purified full-length recombinant WT- and 5mutA-DIMT1 proteins (Supplementary Figure S3A) and performed differential scanning fluorimetry experiments. The results suggest that WT- and 5mutA-DIMT1 are both well-folded proteins and share similar melting temperatures (Figure 3 C and D), as improperly folded, aggregated, or denatured proteins would typically present high background at low temperatures of a melting curve (Gao et al. 2020). Additionally, as shown in Supplementary Figure S3B, the AlphaFold-predicted structure of 5mutA-DIMT1 superimposed very well with the structure of WT-DIMT1 (with a root-mean-square deviation of atomic positions = 0.177 Å). Collectively, these results suggest that 5mutA does not lead to major structural changes in WT-DIMT1.

We then performed electrophoretic mobility shift assays (EMSA) to compare the RNA binding affinities of WT-DIMT1 and 5mutA-DIMT1. We utilized a synthetic RNA probe with carboxyfluorescein at the 5’ end (5’ FAM). This probe bears the same local structure as the two modification sites of DIMT1 in helix 45 of 18*S* rRNA (Supplementary Figure S3C). As shown in Figure 3 E and F, the binding affinity of 5mutA-DIMT1 to the RNA probe significantly decreases in comparison to WT-DIMT1. We determined *K*_half_, which represents the concentration of an enzyme reaching half maximal binding. The *K*_half_ decreases from 1.2 µM protein for WT-DIMT1 to 3.9 µM for 5mutA-DIMT1. Next, we performed *in vitro* methylation assays using WT-DIMT1 or 5mutA-DIMT1 in the presence or absence of SAM. The substrate RNA probe is of the same sequence (but no 5’ FAM) as used in the EMSA assays. The liquid chromatography triple quadrupole mass spectrometry (LC-MS/MS) quantification results show that WT-DIMT1 effectively installs m_2_^6,6^A in this RNA probe while only background level of m_2_^6,6^A can be detected in the reaction using 5mutA-DIMT1 (Supplementary Figure S3 D and E). These results suggest that 5mutA-DIMT1 is weaker than WT-DIMT1 in methylation installation because of the weaker RNA binding ability.

### rRNA binding is a determinant for the nucleolar localization of DIMT1

Since *DIMT1* is an essential gene, we employed a *DIMT1*^+/−^ heterozygous HEK 293T cell line which we established previously (Shen et al. 2020a) to study the cellular localization of WT-, E85A (a catalytically inactive variant), and 5mutA-DIMT1 in cells. We reconstituted empty vector, WT-, E85A- and 5mutA-DIMT1 in this *DIMT1*^+/−^ heterozygous cell line (Shen et al. 2020a). Exogenous Flag-tagged WT-, Flag-tagged E85A-, and Flag-tagged 5mutA-DIMT1 express comparable levels in *DIMT1*^+/−^ heterozygous cells (Supplementary Figure S3F). The immunofluorescence imaging results show that 5mutA-DIMT1 predominantly localizes to the nucleoplasm, while WT- and E85A-DIMT1 both colocalize well with fibrillarin in the nucleolus (Figure 3G). Similarly, MOLM-13C cells expressing WT- or E85A-DIMT1-mClover, but not 5mutA-DIMT1-mClover localize in the nucleolus (Supplementary Figure S3G). The results suggest that rRNA binding may be essential for the primary localization of DIMT1 in the nucleolus, while the enzymatic activity is not required for its localization. DIMT1 does not only interact with 18*S* rRNA, but it also associates with two other ribosome assembly factors (BMS1 and DHX37) as shown in the small subunit processome structure (PDB 7MQ9) (Singh et al. 2021). Immunostaining analyses of MOLM-13C cells which express only exgoneous WT- or 5mutA-DIMT1 shows that BMS1 remains in the nucleolus in both WT- and 5mutA-DIMT1 expressing cells (Supplementary Figure S3H). These data suggest that the RNA-binding deficiency of DIMT1 does not significantly impact the nucleolar localization of its associated proteins.

We further quantified the cellular m_2_^6,6^A levels in 18*S* rRNA using LC-MS/MS. The results showed that the levels of m_2_^6,6^A in 18*S* rRNA are significantly lower in E85A-DIMT1 cells compared to WT-DIMT1, 5mutA-DIMT1, and the empty vector expressing *DIMT1*^+/−^ cells (Supplementary Figure S3I). We previously showed that *DIMT1*^+/−^ heterozygous cells do not have lower m_2_^6,6^A levels in 18*S* rRNA compared to the wild-type HEK 293T cells; however, exogenous E85A-DIMT1 competes with endogenous WT-DIMT1 in binding to 18*S* rRNA and thus leads to lower m_2_^6,6^A levels in 18*S* rRNA (Shen et al. 2020b). In contrast, 5mutA-DIMT1 cells have fully modified 18*S* rRNA, similarly to the empty vector-expressing *DIMT1*^+/−^ cells, suggesting that exogenous 5mutA-DIMT1 cannot compete with endogenous WT-DIMT1 in binding to 18*S* rRNA (Supplementary Figure S3I). These results suggest that the cleft region consisting of the five positive-charged residues are critical for RNA binding and catalysis by DIMT1 *in vitro* and in cells.

### rRNA binding is indispensable for DIMT1 undergoing phase separation

What is the molecular mechanism underlying the nucleoplasmic localization of 5mutA-DIMT1? We first suspected that 5mutA-DIMT1 might disturb the nucleolar-leading sequence (NoLS) in DIMT1. NoLS detector predicted that residues 1 - 26 harbored the nucleolar-leading sequence (Supplementary Figure S4A and B). We fused the predicted NoLS to the N-terminus of mClover. The results show an enrichment of NoLS-mClover in the nucleolus compared to expressing mClover alone (Supplementary Figure S4D). Next, we truncated the first 26 residues and constructed the ΔNoLS-DIMT1. The immunofluorescent imaging reveals ΔNoLS-DIMT1 still remains localized to the nucleolus (Figure S4C). These studies suggest that the *N*-terminal segment of DIMT1 (a putative NoLS sequence) does not play a leading role in DIMT1’s nucleolar localization.

The first 26 residues of DIMT1 are not visible in previous crystal structures despite its presence in the crystallized construct (Shen et al. 2020b), which is likely due to its disordered nature (Supplementary Figure S4E). We used AlphaFold software to predict the structures of WT-DIMT1 (with high per-residue confidence score (pLLDT) over 90% of the amino-acid sequence) (Supplementary Figure S5A). The first 26 residues are predicted to adopt a helical secondary structure with a low pLLDT score below 50% (Supplementary Figure S5B, S5C) and are far from the 5 positive charged cleft residues mutated in this study. Since the nucleolus represents a multilayered biomolecular condensate, whose formation by LLPS facilitates ribosome biogenesis and other functions (Lafontaine et al. 2021), we wondered whether DIMT1 undergoes LLPS and whether this biophysical process is important for its nucleolar localization. Thus, we performed *in vitro* droplet assays using purified DIMT1-mClover (Figure S5D). The results showed that WT-DIMT1 alone does not form droplets *in vitro* using 12.5 µM protein and a series of salt concentrations. Remarkably, the addition of total RNA extracted from HeLa cells promotes the formation of droplets of WT-DIMT1 (Figure 4A). As the RNA concentration increases (16.7 – 100 ng/µL), the apparent volume of the condensed phase (roughly indicated by the total droplet area in each image) increases. The most evident round droplets of WT-DIMT1 with RNA were formed using 200 mM NaCl with 100 ng/µL total RNA since increasing NaCl concentrations to 400 mM and beyond diminished the droplet formation. At identical protein, RNA, and salt concentrations, 5mutA-DIMT1 presents significantly lower, nearly non-detectable droplet formation compared to WT-DIMT1 (Figure 4B). At the optimal condensation formation condition for WT-DIMT1 (200 mM NaCl with 100 ng/µL total RNA), the condensation intensity of 5mutA-DIMT1 is nearly undetectable using the same exposure settings and equal intensity as that for WT-DIMT1 (Figure 5 A and B).

**Fig. 4.**
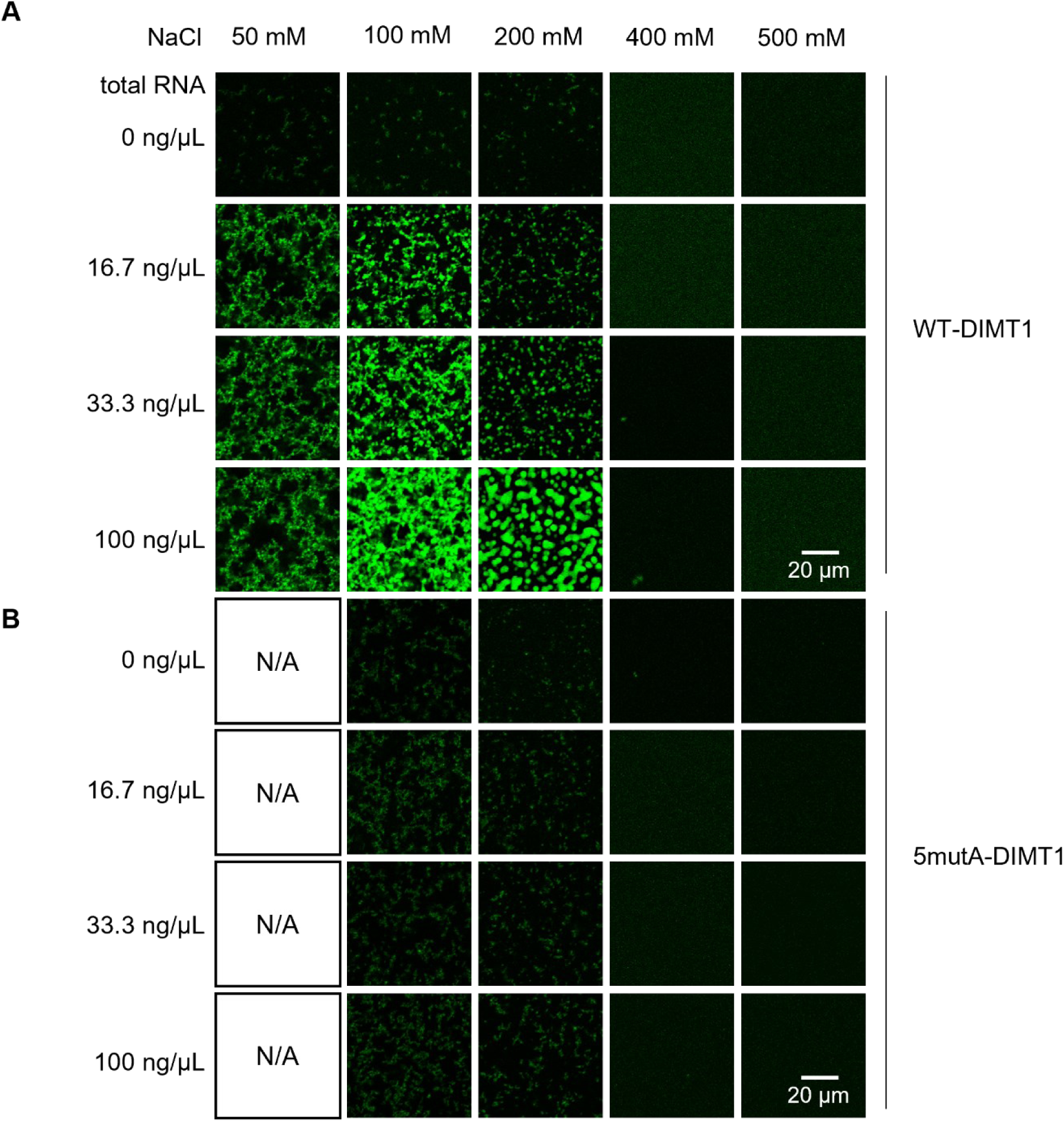
WT-DIMT1 but not 5mutA-DIMT1 forms liquid condensates *in vitro*. *In vitro* droplet formation of 10 µM recombinant (**A**) WT-DIMT1-mClover and (**B**) 5mutA-DIMT1-mClover in the presence of the indicated concentrations of total RNA in the buffer with the indicated salt concentrations. Scale bar, 20 µm.

**Fig. 5.**
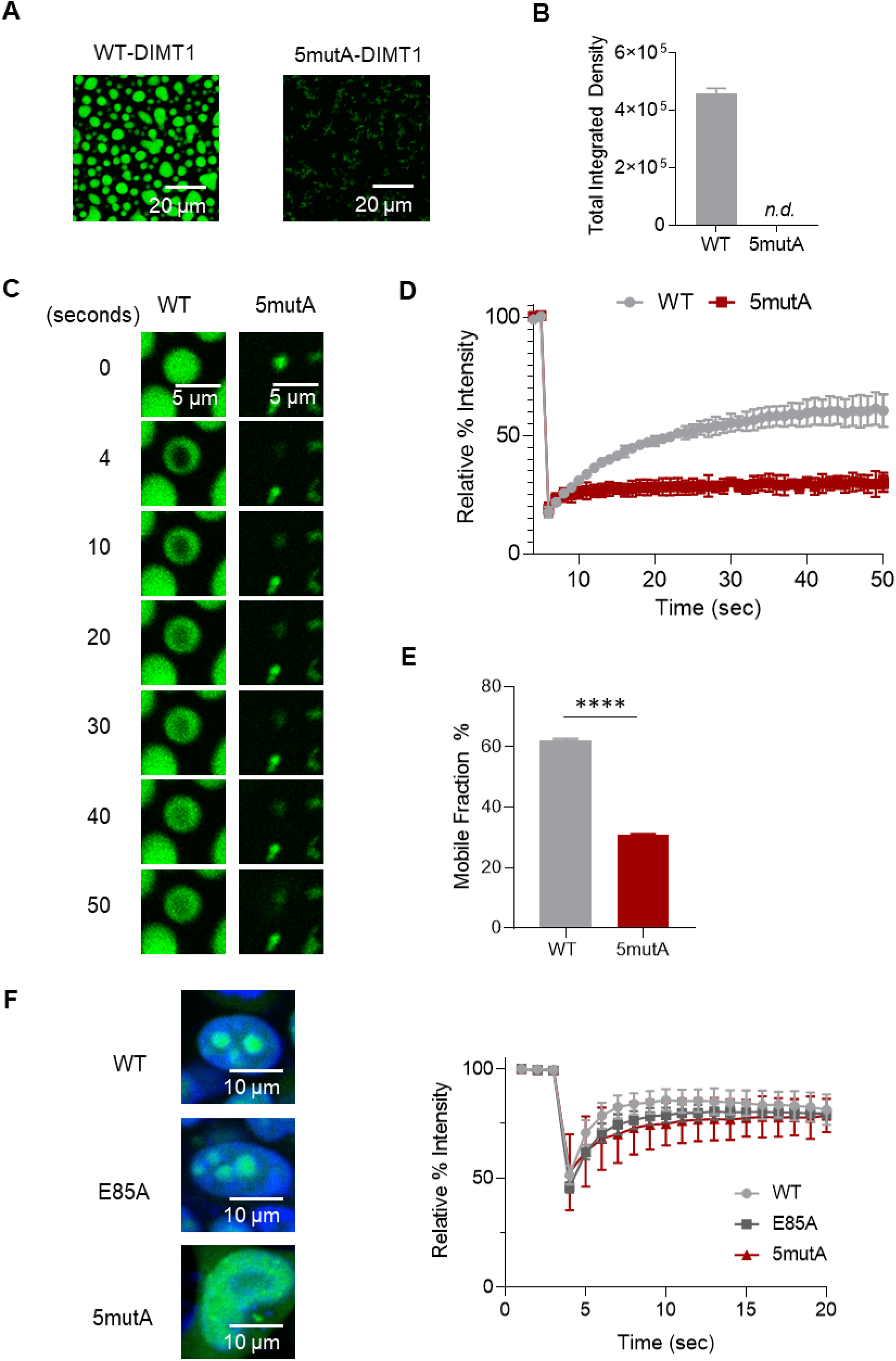
WT-DIMT1, but not 5mutA-DIMT1, forms liquid condensates. (**A**) *In vitro* droplet formation of 10 µM recombinant WT-DIMT1-mClover and 5mutA-DIMT1-mClover in the presence of 100 ng/µL total RNA and 200 mM NaCl. Scale bar, 20 µm. (**B**) Quantification of the total integrated intensity of WT-DIMT1-RNA condensation and 5mutA-DIMT1-RNA condensation in (Fig. 4A). (**C**) Time-lapse images of mClover-WT-DIMT1 (12.5 µM) and 5mutA-DIMT1-mClover (10 µM) droplets from *in vitro* FRAP experiments. Droplets were formed in the presence of 100 ng/µL total RNA and 200 mM NaCl. The FRAP experiments were performed using the same conditions for WT-DIMT1-mClover and 5mutA-DIMT1-mClover. Scale bar 5 µm. (**D**) FRAP curves for the *in vitro* droplets of mClover-WT-DIMT1 (gray) and mClover-5mutA-DIMT1 (maroon). The traces of the FRAP data represent mean ± s.e.m (n = 3, from three independent experiments). (**E**) The mobile fractions from Fig. 5d show that 5mutA-DIMT1 has a significantly slower recovery phase compared to WT-DIMT1. The two-tailed t-test was used to calculate the *p*-value. \*\*\*\**p* < 0.0001. (**F**) Images and FRAP analysis of recovery curves for WT-DIMT1-mClover (gray), E85A-DIMT1-mClover (dark gray), and 5mutA-DIMT1-mClover (maroon) in *DIMT1*^+/−^ HEK 293T cells. The fast fluorescence recovery plots indicate the liquid-like nature of the puncta of WT-DIMT1 and E85A-DIMT1 but not 5mutA-DIMT1. The traces of the FRAP data represent mean ± s.e.m (n = 20 independent experiments, from 3 biologically independent experiments).

We next studied the dynamics of the DIMT1-RNA droplets using fluorescent recovery after photobleaching (FRAP). We measured the recovery halftime and mobile fraction of the DIMT1-RNA droplets by using a one-phase association non-linear regression curve. The recovery halftime for WT-DIMT1-RNA droplets is ∼4.9 seconds, while it is ∼ 9.2 for nucleoplasmic 5mutA-DIMT1 (Figure 5 C and D). We observe that WT-DIMT1 and 5mutA-DIMT1 have mobile fractions of 62% and 30.8%, respectively (Figure 5E). Together, these results suggest that RNA binding is essential for DIMT1 to undergo LLPS *in vitro*.

To assess if DIMT1 undergoes LLPS in the cell nucleolus, we expressed DIMT1 mClover (both WT and E85A) in HEK 293T *DIMT1*^+/−^ cells. As shown in Figure 5F, the mClover-tag on WT and E85A does not disturb the nucleolar localization of the proteins. We then performed FRAP to determine the dynamics of nucleolar WT-DIMT1. FRAP experiments performed on live HEK 293T cells expressing WT-DIMT1 or 5mutA-DIMT1 reveal that the average recovery halftime of WT-DIMT1 nucleolar puncta is 0.8 second, which is noticeably faster than the 1.68 seconds recovery halftime of nucleoplasm 5mutA-DIMT1 (Figure 5F). These results suggest that WT-DIMT1 undergoes LLPS *in vitro* and in cells, and rRNA binding-promoted LLPS of WT-DIMT1 is a possible driving force for the nucleolar localization of DIMT1.

### The non-catalytic rRNA binding ability of DIMT1 is critical for cell proliferation

To assess the significance of the nucleolar localization of DIMT1 in promoting cell proliferation, we performed competition-based proliferation assays using CRISPR-resistant WT-, 5mutA-, and ΔNoLS-DIMT1 in MOLM-13C cells. The western blot analyses suggest that 5mutA-DIMT1 was expressed close to the endogenous level of DIMT1 in MOLM13C cells (Supplementary Figure S5A). As shown in Figure 6A, MOLM-13C cells re-expressing 5mutA-DIMT1 showed the same proliferation as cells re-expressing an empty vector, both presenting significant decreases in cell proliferation compared to cells re-expressing wild-type DIMT1. In contrast, WT- and ΔNoLS-DIMT1-expressing cells do not present decreased cell proliferation after transduction of the sgDIMT1 targeting endogenous DIMT1 (Figure 6A). Pre-rRNA processing in MOLM-13C cells expressing 5mutA-, WT-, E85A-DIMT1 after transduction with sgRosa or sgDIMT1 was analyzed by using Northern blotting (Figure 6B). Depletion of endogenous DIMT1 caused significant accumulation of 21*S* precursor only in 5mutA-DIMT1, but not in WT-, and E85A-DIMT1 expressing cells. These results indicate that only WT- and E85A DIMT1, but not the RNA-binding deficient 5mutA-DIMT1 are able support 18*S* rRNA processing. Polysome profiling of MOLM-13C cells expressing 5mutA-, WT-, E85A-DIMT1 with the endogenous DIMT1 depleted demonstrated decreased levels of polysomes only in 5mutA-DIMT1 cells (Figure 6C). To determine the translational status of the top five mRNA targets identified by ribo-seq in DIMT1 depletion compared to the control, we performed qRT-PCR on RNA isolated from the light and heavy fractions of the polysome profiles. The results showed that the five mRNAs which presented increased ribosome occupancies upon DIMT1 depletion in ribo-seq are enriched in the heavy fractions of the gradients in cells expressing 5mutA-, but not WT- or E85A-DIMT1 (Figure 6D). Altogether, these results consistently demonstrate that RNA binding-facilitated nucleolar localization but not the methyltransferase activity of DIMT1 is essential for ribosome biogenesis to support AML proliferation.

**Fig. 6.**
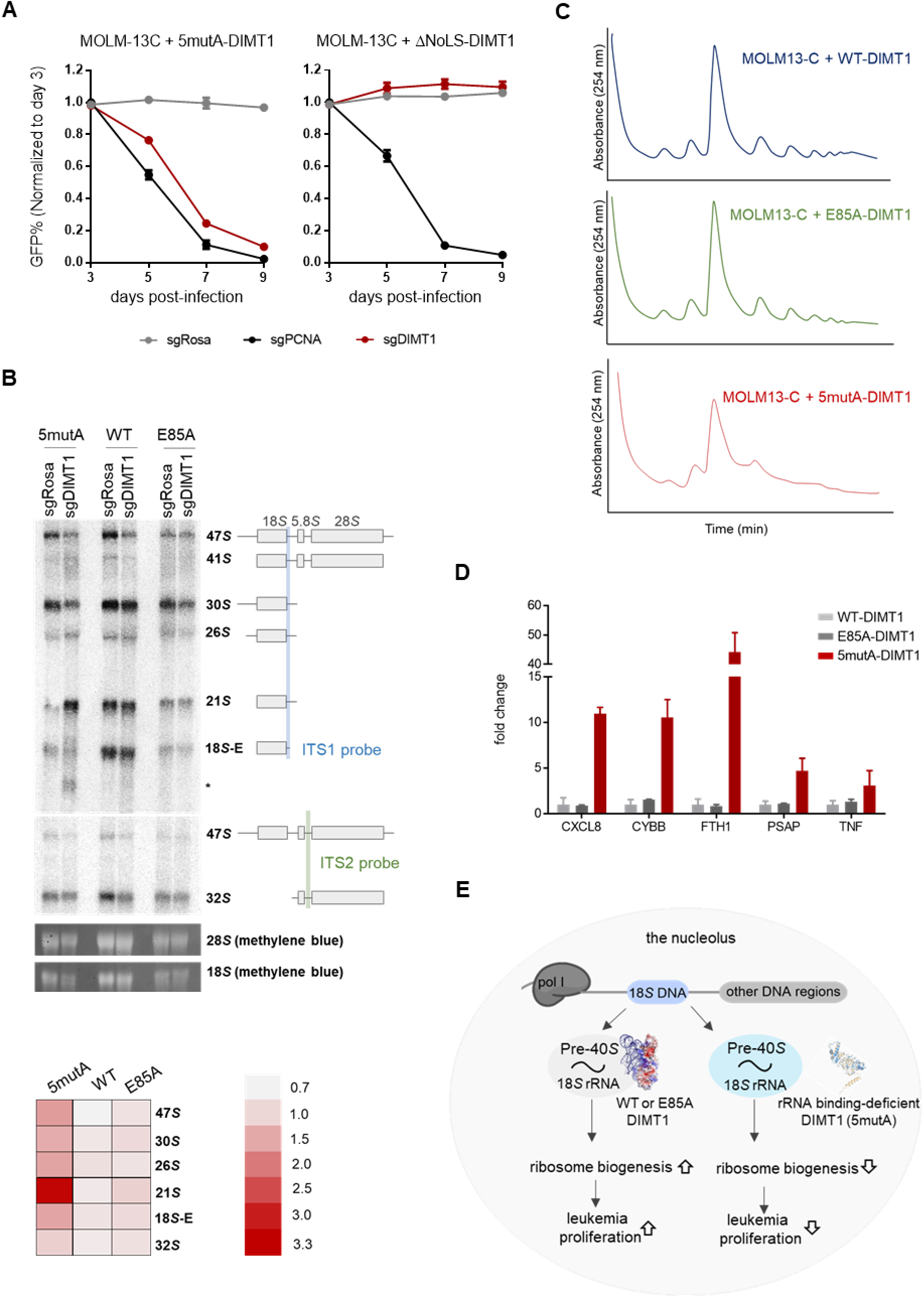
The RNA binding region but not the catalytic activity of DIMT1 is required for cell proliferation. (**A**) Competition-based proliferation assay of MOLM-13C cells re-expressing sgRNA-resistant 5mutA-DIMT1 (left), and ΔNoLS-DIMT1 (right). Cells were transduced with the indicated sgRNAs (sgRosa as a negative control, sgPCNA as a positive control, and sgDIMT1). (**B**) Upper panel: representative Northern blots showing pre-rRNA processing in MOLM-13C cells re-expressing sgRNA-resistant 5mutA-, WT-, and E85A-DIMT1 after transduction with the indicated sgRNAs. Northern blot analyses were performed using two probes targeting ITS1 and ITS2. The detected pre-rRNA species are indicated to the right and schematized. Lower panel: Abundance of rRNA precursors of two biological replicates were quantified with Phosphorimager and converted into a heatmap. The heatmap shows the abundance of each pre-rRNA species in sgDIMT1 infected cells compared to the abundance of the same pre-rRNAs in cells infected with sgRosa. (**C**) Polysome profiles of MOLM-13C cells expressing exogenous WT/E85A/5mutA-DIMT1 after knockout of endogenous DIMT1. (**D**) Results of the qRT-PCR of RNA extracted from the light or heavy fractions of the polysome profiles shown in C performed in 2 biological replicates. Y axis represents the abundance of mRNA in the heavy fraction relative to those in the light fraction. The fold change values of the relative mRNA abundance in WT-DIMT1 cells were normalized to 1. (**E**) Schematic illustration of a positively charged cleft in DIMT1 supports 40*S* small subunit assembly and leukemia proliferation. The hollow arrows indicate up (stimulating) or down (inhibiting) regulation of the proliferation.

## Discussion

Previous studies have emphasized that dysregulation of ribosome assembly results in distinct types of human diseases, including cancers. However, there is limited knowledge about the underlying mechanism by which ribosome assembly factors and enzymes lead to these diseases. Here we investigated non-catalytic role of ribosome assembly factor DIMT1 in cell proliferation. Results of competition-based proliferation assays demonstrated that AML cell lines expose higher sensitivity to DIMT1 loss compared to the tested solid tumor cell lines. Ribosome profiling of AML cells upon DIMT1 depletion revealed subsets of dysregulated transcripts, including MYC and HOX mRNAs implicated in variety of different cancers, suggesting that DIMT1 affects common cancer pathways and may impact a broad spectrum of cancers including leukemia.

Using catalytically dead DIMT1 mutant we showed that MOLM-13C cells rely on DIMT1 not for its catalytic activity but for its ability to bind to 18*S* RNA for its nucleolar localization and function (Figure 6C). Of note, our study of 5mutA- and ΔNoLS-DIMT1 reveals that the rRNA binding but not the predicted nucleolar-leading sequence is the driving force for nucleolar localization. Taken together, our study on a positively charged cleft in DIMT1 provides insight into the vital role that rRNA binding plays in the nucleolar phase separation and functions of DIMT1. This discovery opens new avenues to studying other ribosome assembly factors and their connections to cancers.

### rRNA-binding affinity but not the putative nucleolar localization sequence is important for the nucleolar localization of DIMT1

The nucleolus is the hallmark of nuclear compartmentalization and plays a primary role in rRNA synthesis and ribosome assembly. DIMT1 localizes in the nucleolar compartment. Previous studies show that short protein sequences with a minimum of five basic amino acids (often lysine and arginine residues) are termed nucleolar localization sequences (NoLS) (Martin et al. 2015). Nuclear proteins without nucleolar function are often less concentrated or completely excluded from the nucleoli, while nucleolar proteins are highly enriched in this nuclear compartment. Although NoLS have been identified in nucleolar proteins, it remains unclear which properties or requirements of such proteins are essential for accumulation in the nucleoli. We noticed that DIMT1 contains a predicted NoLS at the *N*-terminus of the protein (Supplementary Figure S4A). However, deleting this region did not alter the nucleolar localization of DIMT1. In contrast, mutations of a basic RNA binding cleft which do not disturb the predicted NoLS are responsible for modulating the nucleolar localization of DIMT1, which highlights the role of rRNA-binding in cellular localization.

### RNA-promoted phase separation is a possible mechanism for DIMT1 localization in the nucleolus

The nucleolus is the most prominent nuclear compartment and is formed by liquid-liquid phase separation of proteins and protein/RNA complexes. This multilayered sub-cellular compartment performs a host of diverse roles in a highly organized manner. In this study, we investigated whether DIMT1 undergoes liquid-liquid phase separation *in vitro* and inside cells. Our results suggest that RNA binding promotes *in vitro* droplet formation of WT but not 5mutA-DIMT1. Additionally, WT-DIMT1 presents a liquid condensation property in the nucleolus. These results suggest that RNA-promoted LLPS is a possible mechanism for DIMT1 localizing in the nucleolus. However, this study does not exclude the possibility that 18*S* rRNA-binding facilitates the nucleolar localization of DIMT1 regardless of its LLPS status.

The localization of proteins and RNAs to the nucleolus is critical for a host of biological events (Lafontaine et al. 2021). The localization or exclusion of proteins from the nucleolus can dramatically change in response to cellular stimuli and reorganization during cell cycle progression. DIMT1 is an indispensable assembly factor in 18*S* rRNA processing and 40*S* subunit assembly. Thus, the function of DIMT1 may better be facilitated by its high concentration in the nucleolus. Additionally, DIMT1-mediated m_2_^6,6^A occurs at two adjacent sites in 18*S* rRNA at nearly 100% occupancy. We predict that enriched DIMT1 in the nucleolus might be indispensable for its catalysis. Although DIMT1-mediated m_2_^6,6^A is not required for the proliferation of MOLM-13C cells, the double methylation does play a role in other cellular aspects such as cell differentiation, especially since the loss of DIMT1-mediated m_2_^6,6^A sensitizes living organisms to stress conditions and decreases translational fidelity (Helser et al. 1971; Lafontaine et al. 1994; Tokuhisa et al. 1998; Wieckowski and Schiefelbein 2012).

### The rRNA cleft identified in this study is an Achilles’ heel of DIMT1

DIMT1 is a common essential gene that is indispensable for ribosome biogenesis. However, our results suggest that certain cell types such as AML cells are more sensitive to DIMT1 deficiency (Figure 1A). In our previous study (Shen et al. 2021), we observed that *DIMT1*^+/−^ HEK 293T expressing E85A-DIMT1 has reduced translation and cell proliferation compared to *DIMT1*^+/−^ HEK 293T expressing the WT-DIMT1. But E85A DIMT1-expressing MOLM-13C cells have indistinguishable cell proliferation from WT-DIMT1 cells. It is not known whether E85A DIMT1 MOLM-13C cells have comparable translational status to WT DIMT1 MOLM-13C cells. We reasoned that the cell type specificity might contribute to the different requirements of DIMT1’s catalytic role in the cell proliferation of HEK 293T and MOLM-13C. Additionally, the *DIMT1*^+/−^ HEK 293T system still contains remaining endogenous DIMT1, which is different than the MOLM-13C system which expresses E85A-DIMT1 before complete ablation of endogenous DIMT1.

Since we show that DIMT1 supports MOLM-13C cell proliferation through promoting rRNA processing and ribosome biogenesis, methods to regulate DIMT1’s non-catalytic role in ribosome assembly could be further pursued. Given that the data in this study showed the RNA-binding cleft is linked to DIMT1’s nucleolar localization and function (Figures 3G and 6), it may be a new therapeutic site for targeting DIMT1 in rapidly dividing cells, such as in AML proliferation. Identifying this indispensable role of an rRNA binding cleft remote from DIMT1’s catalytic site is important and potentially inspiring because there are several other rRNA modifying enzymes and rRNA binding proteins that also function as assembly factors in pre-ribosomal complexes in the nucleolus. Tackling the RNA binding ability of these similar rRNA modifying enzymes and rRNA binding proteins may provide a new therapeutic avenue for ribosomopathies.

In summary, this study describes an RNA binding cleft of DIMT1 and uncovers its function in controlling DIMT1’s cellular localization and role in promoting cell proliferation. The findings detailed in this report open new avenues for studying ribosome assembly factors and introduce a potential therapeutic target in treating ribosome defects such as those in AML and other human disorders related to ribosomal defects.

## Materials and Methods

### Mammalian cell culture, lentivirus production and transduction

HEK293T, HUH7C, and A375C cells were cultured in DMEM + GlutaMAX (GIBCO, 11995065) with 10% FBS (GIBCO, 26140-079) and 1% Pen/Strep (Corning, 30-002-CI) in a humidified incubator with 5% CO_2_ at 37°C. MOLM-13C and MV4-11C cells were cultured in RPMI (GIBCO, 22400-089) supplemented with 10% FCS and 1% Pen/Strep.

For lentivirus production, HEK293T cells in a 10-cm plate at 90-100 % confluency were transfected with 10 μg plasmids of interests, 5 μg VSV-G plasmid, and 7.5 μg psPAX2 plasmid using 80 µL of 1mg/mL polyethylenimine (Polysciences, 25000) in 500 μL of Opti-MEM (Gibco, 31985062). The media was changed with 6 mL fresh DMEM 6 - 8 hours after transfection. Lentivirus was collected twice a day for the next three days after transfection. Collected virus was pooled together, filtered with a 0.45 μm PVDF filter (Millipore) and stored at −80 °C in small aliquots for long-term use.

For viral transduction, cells of interest were transduced with lentivirus using and 4 μg/mL polybrene (Sigma, H9268) and centrifuged at 650 × *g* for 25 minutes at room temperature. The transduced cells were incubated overnight at 37 °C, and media was replaced 15 hours post-infection.

### Construction of stable cell lines

HEK293T stable cell lines were generated in HEK293T *DIMT1*^+/−^ heterozygous background using pPB vector as previously described (Shen et al. 2020b). Transfection was performed at 60% cell confluency using Lipofectamine 2000 (Invitrogen, 11668019). 24 hours after transfection, the cells were selected under 2 μg/mL puromycin for 2 weeks. During the selection period, cells were resuspended every 2 days with fresh DMEM sufpplemented with 10% FBS and 2 μg/mL puromycin. The stable overexpression was confirmed by Western blotting using an anti-FLAG antibody (Thermo, MA1-91878-HRP).

To create MOLM-13C stable cell lines, MOLM-13C cells were transduced with WT- and E85A-CRISPR-resistant DIMT1 followed by a P2A-linked mCherry. Transduced MOLM-13C cells containing roughly 30% mCherry positive cells were sorted and collected using a FACS MoFlo Astrios Cell sorter (Beckman).

### Plasmid constructs

For DIMT1 and recombinant DIMT1-mClover protein expression: DIMT1 mutations were introduced through site-directed mutagenesis PCR into a pET-28a vector to generate DIMT1 5mutA. DIMT1-mClover constructs were introduced into pET-28a through fusion PCR to generate a DIMT1-mClover inserts. This was used to generate DIMT1 WT pET-28a-mClover, DIMT1 E85A pET-28a-mClover DIMT1, and 5mutA pET-28a-mClover vectors.

For transfection and establishment of HEK 293T stable cell lines: DIMT1 5mutA pPB vector was generated by subcloning 5mutA insert from existing DIMT1 5mutA pET-28a vector. Similarly, DIMT1 mClover constructs were subcloned out of pET-28a-mClover vectors into pPB vectors to generate DIMT1 pPB-mClover vectors. The ΔNoLS was generated by truncating amino acids 1-26 and inserting into pPB and pPB-mClover vectors.

To generate sgRNA encoding plasmids, sense and antisense DNA oligos were annealed and ligated into a BsmbI digested LRG2.1 expressing GFP (Addgene: 108098).

To generate CRISPR-resistant DIMT1 mutant, the following base substitutions were introduced into the DIMT1 cDNA: A177->T, G180->T; G183->T; A186->G, T189->A by PCR mutagenesis. CRISPR-resistant DIMT1 was cloned into Lenti Cherry vector using In-Fusion cloning system (Takarabio, 638943). Generated vectors encode N-terminus Flag-tagged CRISPR-resistant WT- and E85A-DIMT1, which are followed by P2A-linked mCherry.

All plasmids were validated through sanger sequencing.

### Protein purification

pET-28a vectors were transformed into *Escherichia coli* BL21(DE3, C2527H) for purification. Bacteria were grown at 37 °C until an optical absorbance at 600 nm between 0.6 – 0.8 was achieved, at which point protein expression was induced by the addition of IPTG at 1 mM final concentration. After overnight growth at 16 °C, cells were collected and lysed using lysis buffer (25 mM Tris-HCl, pH 7.5, and 500 mM NaCl). Cell lysate was centrifuged at 4 °C at 11,000 rpm for 30 min and the supernatant was then loaded onto a Ni-affinity chromatography column (GE Healthcare), which was pre-equilibrated using the lysis buffer. The loaded column was washed with 100 mL wash buffer (50 mM imidazole in lysis buffer) before the bound protein was eluted with elution buffer (500 mM imidazole in lysis buffer). Finally, eluted protein was further purified by running a HiLoadSuperdex 200 (GEHealthcare, 28989335) column with S200 buffer (25 mM Tris-HCl, pH 7.5, 500 mM NaCl, 5% Glycerol).

### Differential scanning fluorimetry assay

Purified WT-DIMT1 and 5mutA-DIMT1 proteins were diluted to 1 mg/mL in S200 buffer (25 mM Tris-HCl, pH 7.5, 500 mM NaCl, 5% Glycerol). 19 μL of each protein was transferred to one well in a 384 well plate, and 1 μL of 5-fold SYPRO Orange (Thermo Fisher, S6650) was added to each well. The plate was sealed and spun at 3,600 *g* for 2 min. The fluorescent signal at 570 nm was recorded using a RT-qPCR machine with the temperature ramping from 20 to 95 °C. The data were analyzed using DSF World (Wu et al. 2020).

### Competition assay

To perform proliferation-based cellular competition assay, Cas9-expressing human cell lines of interest were transduced with sgRNA co-expressed with GFP as a marker for transduction. To deplete DIMT1, sgDIMT1 # 1 and sgDIMT1 # 2 were used, while sgRosa and sgPCNA were used as negative and positive controls, respectively, (sequences were listed in the Supplementary Table 1). Number of GFP-positive cells was measured every second day starting from day 3 post-infection using Guava Easy 5HT instrument (Luminex, 0500-4005). Percentage of GFP-positive cells on each day was normalized to percentage of GFP-positive cells on day 3 post-infection. The competition assays were performed in three biological replicates, each in two technical replicates.

### EMSA

A FAM labeled RNA probe, 5’-rUrUrCrCrGrUrArGrGrUrGrArArCrCrUrGrCrGrGrArA-3’, was dissolved in RNase-free water at 100 μM and diluted to 1 μM in EMSA binding buffer (25 mM Tris-HCl, pH 7.2, 150 mM NaCl, and 40 U/mL RNasin). WT- and 5mutA-DIMT1 were diluted to a concentration series of 0.1 μM, 0.5 μM, 1 μM, 2 μM, 4 μM and 6 μM in EMSA binding buffer. 1 µL RNA probe and 1μL protein were mixed with 8 μL binding buffer and incubated on ice for 30 min. The entire 10 μL RNA–protein mixture was then loaded onto a 4% TBE gel with 2.5 μL loading buffer (1% bromophenol blue and 50% glycerol) and run for 35 min at 120 V in a 4 °C. The gel was imaged using a Amersham^TM^ Typhoon^TM^ 5 (cytiva, 29187191) fluorescent scan with 473 nm laser. Quantification was carried out using Fiji ImageJ to quantify intensity of the bottom free-RNA band. The *K*_D_ (dissociation constant) was calculated in Prism with nonlinear curve fitting (allosteric sigmoidal) with equation y = V_max_*x^h^/(K_half_^h^ + x^h^) where y is the bound fraction, x is the concentration of DIMT1 (μM), V_max_ is set to 1, h is the hill slope, and *K*_half_ is the DIMT1 concentration resulting in 0.5 bound fraction.

### RNA isolation

Total RNA was extracted from cell pellets using TRIzol reagent (Invitrogen,15596026) following manufacturer’s instructions. To isolate 18*S* rRNA isolation, 5 μg total RNA was denatured in RNA loading dye (Thermofisher, R0641) for 10 min at 72 °C and ran on a 1% TBE agarose gel. The 18S rRNA band was then excised from the gel and purified using a gel RNA recovery kit (Zymoclean, R1011).

### Quantitative analysis of the m_2_^6,6^A level

Purified RNA was digested and dephosphorylated to single nucleosides using nucleoside digestion mix (NEB, M0649S) at 37 °C for 2 hours. The nucleosides were quantified using retention time and nucleoside-to-base ion mass transitions of 268.0 → 136.0 (A), 284.0 → 152.0 (G), and 296.0 → 164.1 (m_2_^6,6^A). The percentage ratio of m_2_^6,6^A to A was then used to compare the different modification levels between samples. LC-MS/MS data was analyzed using Skyline software.

### *In vitro* methylation assay

The *in vitro* methylation assays were performed in a 30 mL reaction mixture containing the following components: 4 µg biotinylated RNA probe 3’-rUrArGrGrUrGrArArCrCrUrG-5’ (0.945 nmol), 0.7875 nmol protein, 1 mM SAM, 50 mM Tris-HCl pH 7.5, 5 mM MgCl_2_, and 1 mM DTT. The reaction mixture was incubated at 16 °C for 10 hours. After incubation, streptavidin beads (Thermo Scientific, 10608D) were used to purify the RNA probe, following the instructions from the manufacturer, and eluted with 18.5 μL RNase-free water at 75 °C for 5 min.

### Immunofluorescence microscopy

*DIMT1*^+/−^ HEK 293T cells expressing exogenous DIMT1 were grown in a 6 well plate on top of cover slide. Cells were rinsed once with 1 × PBS followed by a 15-minute rocking incubation with 1 mL of 4% Formaldehyde in PBST (1 × PBS, 0.05% Tween-20). Next, cells were incubated in 0.5% Triton in 1 × PBST for 20 minutes followed by rinsing with PBST. Cells were then blocked in 1% BSA in PBST for 1 hour followed by 1 hour in primary anti-FLAG antibody (1:1000, MA1-142) incubation in 1% BSA in PBST. Cells were then rinsed with 1% BSA in PBST followed by a 1-hour incubation in anti-Rat secondary antibody (1:300, A-11006). Following secondary incubation, cells were rinsed with PBST followed by 1 minute incubation with 0.5 μg/mL DAPI and another PBST rinse. 20 μL Anti-fade reagent (P36970) was then applied to a microscope slide and the cover slip sealed on top. Microscope slides were then imaged on a Leica TCS SP8 confocal microscope.

### Fluorescence recovery after photobleaching (FRAP)

All FRAP assays were performed using the bleaching module of the Zeiss LSM 880 confocal microscope. The 488 nm laser was used to bleach the mClover signal. Bleaching was focused on a circular region of interest (ROI) using 100% laser power, and time-lapse images were collected afterward. Additionally, a circular area of the same size outside the bleaching point was used as an unbleached control. The fluorescence intensity was directly measured in the Zen software (Zeiss). The values are reported as relative to pre-bleaching time points. GraphPad Prism was used to plot the data. The halftime for each replicate was calculated using a non-linear one-phase association fit with equation of Y = Y_0_ + (Plateau - Y_0_) * (1 - exp(-K*x)) in which Plateau-Y_0_ is the slow recovery fraction, K is the recovery rate, and the half time is ln(2)/K. The mobile fraction is represented as the Plateau.

To perform *In vitro* FRAP experiments, 30 µL *in vitro* LLPS reaction was set up at room temperature with 25 mM Tris-HCl and NaCl concentrations of 50 mM, 100 mM, 200 mM, 400 mM, and 500 mM, DIMT1-mClover at a 12.5 μM, and total RNA at 0 ng/μL, 16.7 ng/μL, 33.3 ng/μL, and 100 ng/μL. Reactions incubated at RT for 20 minutes and transferred to a 384-well glass bottomed reader plate and spun at 100 × g for 1 minute. Images were taken using a Zeiss LSM 880 confocal microscope under 20× lens.

The cellular FRAP was performed on HEK293T cells expressing DIMT1-mClover constructs cultured in glass bottom dishes (MatTek, P35GC). The media was changed with FluoroBrite DMEM Media (Thermo Scientific, A1896701) and cells were incubated with 1 drop of NucBlue™ Live ReadyProbes™ Reagent (Thermo Scientific, R37605) for 1 minute before imaging with a Zeiss LSM 880 confocal microscope under 20 × lens.

### Protein quantitation and Western blotting

Protein concentrations were determined by Bradford assay (5000006, Bio-Rad). 50 μg of protein samples were boiled at 95 °C in 1 × Laemmli sample buffer for 5 minutes followed by loading onto SDS-PAGE gel and running at 180 V for 1 hour. Samples were transferred onto PVDF membranes using a semidry transfer apparatus at 20 V (sequences were listed in the Supplementary Table 1) for 50min. Membranes were then blocked with 3% milk in PBST followed by primary antibody incubation in 3% milk PBST. Membranes were then rinsed with PBST followed by secondary incubation. The membranes were then washed with PBST and visualized using an ECL Western blotting detection kit (Thermo Fisher, 34577).

### AlphaFold structure analysis

The DIMT1-5mutA amino acid sequence was input into AlphaFold’s ColabFold program. The resulting output PyMol file was compared to the solved crystal structure of WT-DIMT1 and the AlphaFold protein structure database model of WT-DIMT1. PyMol’s superimpose command was then used to determine how closely folded the different structures were.

### Ribosome profiling

MOLM-13C were transduced with sgDIMT or sgRosa co-expressed with GFP as a transduction marker in three biological replicates. Efficiency of lentivirus transduction was estimated by counting number of GFP-positive cells using Guava Easy 5HT instrument (Luminex, 0500-4005). Depletion of DIMT1 was confirmed by Western blotting. On day 5 post-infection, cells were treated with 100 μg/mL cycloheximide (CHX) for 7 min. The cells were harvested by spinning at 400 *g* for 5 min and then washed with ice-cold PBS supplemented with CHX (100 μg/mL) once.

The collected cell pellets were resuspended in lysis buffer (10 mM Tris-HCl at pH 7.4, 150 mM KCl, 5 mM MgCl_2_, 100 mg/mL CHX, 0.5% Triton X-100, Protease Inhibitor Cocktail, Rnase inhibitor, and 100 μg/mL CHX) and kept on ice for 15 min, followed by centrifugation at 13,000 rpm for 15 min at 4°C. For the input library construction, 10 μL of 10% SDS was added to 100 μL of the cell lysis aliquot, and the Zymo Research RNA Clean and Concentrator-25 kit was used to purify total RNA. On the other hand, to digest mRNA not protected by ribosome, 120 U Rnase I (Ambion) the cell lysate (Absorbance_260 nm_ = 3) of and incubated on ice for 30 min. The reaction was stopped by addition of SUPERase:In (Ambion). To obtain ribosome protected fragments (RPF), the digested lysate was loaded on top of 10 - 50% w/v sucrose gradient prepared in the lysis buffer without Triton X-100 by the Gradient Master (BioCamp). The gradients were centrifuged at 38,300 rpm for 2 hours and 45 min at 4 °C in a SW-40Ti (Beckman) rotor using an Optima XE-100 Ultracentrifuge (Beckman). Then, the gradient was fractionated into 1 mL by a Biocomp piston fractionator with a TRIAX flow cell for continuous UV detection at 256 nm, and the monosome fractions were pooled, followed by RNA purification with The Zymo Research RNA Clean and Concentrator-25 kit. Obtained RNA was depleted of rRNA with Ribo-minus kit (Invitrogen), and then was separated in 15% TBE-Urea gel and stained with ethidium bromide. Gel slices containing nucleic acid 27 - 30 nucleotides (nts) were excised and purified using Zymo small RNA gel purification kit. To prepare the input samples, RNA longer than 200 nts were extracted from the total RNA using RNA extraction kit (Zymoresearch, R1013). Purified RNA was then fragmented using RNA Fragmentation Reagents kit (Invitrogen). Prior library construction, both input sample RNA and RPF RNAs were subjected to RNA end repair under PNK treatment (Thermofisher). Libraries were constructed according to NEBNext Small RNA Library Prep kit (NEB) protocol. The concentrations for all the libraries were determined by KAPA Library Quantification Kit (KAPA Biosystems) following the manufacturer’s protocol and subjected to Next-generation high-throughput sequencing using an illumina NextSeq 550 with a single-end 75-bp read length.

### Data analysis

The high throughput sequencing reads were subjected to adaptor trimming using *Trimmomatic* (Bolger et al. 2014) with following command: trimmomatc SE -phred33 ILLUMINACLIP:TruSeq3-SE.fa:2:30:10 MAXINFO:20:0.5 MINLEN:20. Reads mapping to rRNA, tRNA and mtDNA were firstly cleaned using bowtie2. The rRNA sequences were download from the genebank. The tRNA seqeunces were downloaded from gtRNA database. The mtDNA seqeunce were directly obtained from the whole genome sequence. Then, the clean reads were mapped to genome GRCh38 using *STAR* (Dobin et al. 2013) with following command: STAR -- runThreadN 8 --alignSJDBoverhangMin 1 --alignSJoverhangMin 51 --outFilterMismatchNmax 2 --alignEndsType EndToEnd --readFilesCommand gunzip -c --outFileNamePrefix --quantMode GeneCounts --outSAMtype BAM SortedByCoordinate --limitBAMsortRAM 31532137230 -- outSAMattributes All. Lastly, *featureCounts* (Liao et al. 2014) was used to count the read numbers for each gene. The differential translation efficiency and gene expression were calculated using DESeq2 (Love et al. 2014) followed the instruction from a previous reported procedure (Chothani et al. 2019).

### Northern blots

10 μg of total RNA were resolved on a denaturing 1.25 % agarose gel containing 6.7% formaldehyde. The RNA was transferred overnight onto a Hybond N membrane by passive transfer in 20× saline–sodium citrate. After UV crosslink, RNA was probed for pre-rRNAs using P^32^-labeled oligonucleotides targeting ITS1 (CCTCGCCCTCCGGGCTCCGTTAATGATC) or ITS2 (GGGGCGATTGATCGGCAAGCGACGCTC). Membranes were hybridized (overnight at 65°C), washed and dried prior to exposure and signal detection in a Typhoon FLA 9500 (GE Healthcare, Chicago, IL, USA) phosphoimager. The pre-rRNAs were quantified and normalized to the mature 28*S* rRNA (EtBr or methylene blue staining) and then to the control samples (sgRosa).

### Polysome profiling followed by qRT-PCR

MOLM-13C cells expressing WT-, E85A-, or 5mutA-DIMT1 on day 5 post-infection with sgDIMT1 were treated with 100 μg/mL cycloheximide (CHX) for 7 min. The cells were harvested by spinning at 400 g for 5 min and then washed with ice-cold PBS supplemented with CHX (100 μg/mL). The collected cell pellets were resuspended in lysis buffer (10 mM Tris-HCl at pH 7.4, 150 mM KCl, 5 mM MgCl2, 100 mg/mL CHX, 0.5% Triton X-100, Protease Inhibitor Cocktail, Rnase inhibitor, and 100 μg/mL CHX) and kept on ice for 15 min, followed by centrifugation at 13,000 rpm for 15 min at 4°C. The lysates were loaded on top of 10 - 50% w/v sucrose gradient prepared in the lysis buffer without Triton X-100 by the Gradient Master (BioCamp). The gradients were centrifuged at 38,300 rpm for 2 hours and 45 min at 4 °C in a SW-40Ti (Beckman) rotor using an Optima XE-100 Ultracentrifuge (Beckman). Then, the gradients were fractionated into 1 mL by a Biocomp piston fractionator with a TRIAX flow cell for continuous UV detection at 256 nm. After fractionation 200 μl of the light fraction (fraction1) and 200 μl of the heavy fractions (pooled fraction 5-12) were subjected to RNA extraction. To have a normalization control, 0.2 ng of Luciferase mRNA were spiked-in prior each RNA extraction. Extracted RNA from light and heavy fractions were subjected to qRT-PCR using Luna RT-qPCR kit (NEB).

## Acknowledgments

This work was supported by the National Institutes of Health R35 GM133721, R01 HL160726, RSG-22-064-01-RMC American Cancer Research Scholar, and Damon Runyon Innovator Award to KF. Liu, R35 GM118090 to R. Marmorstein, R01 CA258904 to JW. Shi, T32GM07133398 to K. Schultz, and Heinrich Kronstein Foundation to Y. Gonskikh. We thank Dr. Matthew Kayser for sharing the Leica SP8 confocal microscope and Dr. Brain Capell for sharing the Nextseq550 sequencer.

## Data availability

The sequence data have been deposited in the NCBI GEO database under the accession code GSE197276. All other data are available from the corresponding author upon reasonable request. All the primers and probes used in the paper are listed in Table S3.

## Author contribution

J.S., Y.G. H.S. and K.F.L. designed the experiments. Y.G. prepared the ribo-seq samples. Y.G., K.B., and B.P. performed the competition-based proliferation assays with the guidance from JW.S. H.S., processed the ribo-seq data. J.S., Y.G., and H.S., generated the constructs and the mammalian cell lines used in this study. J.S., Y.G., and H.S. purified the recombinant DIMT1 (full-length proteins and the truncation variants) used in this study. J.S. performed the immunofluorescence imaging experiments and analyzed the data. J.S., Y.G., and H.S. wrote the manuscript together with K.F.L. All authors participated in the discussion and editing of the manuscript.

