## Supplemental Text and Figures for "Non-catalytic regulation of 18*S* rRNA methyltransferase DIMT1 in acute myeloid leukemia"

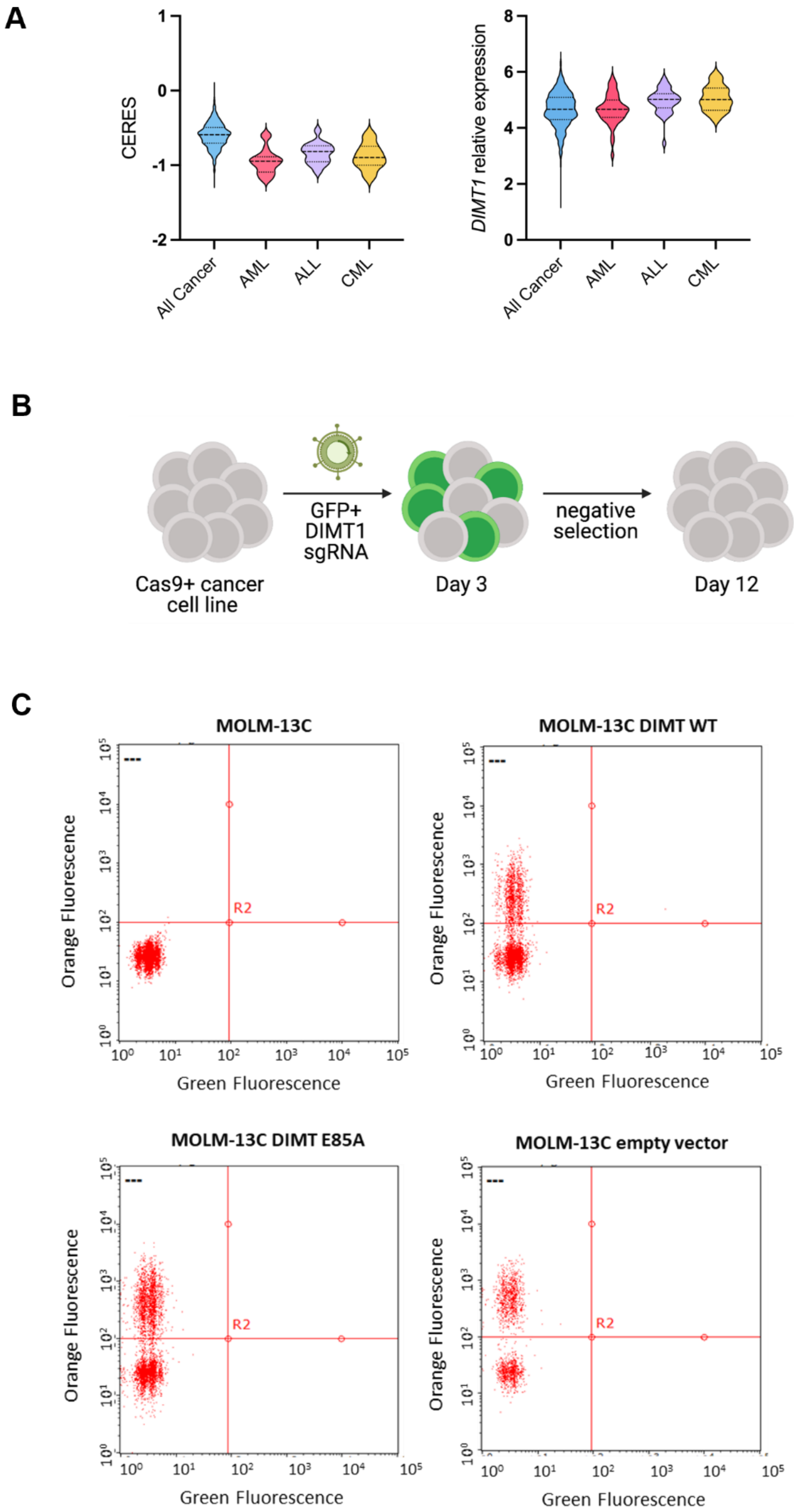
A
CERES
1
0
-1
-2
All Cancer
AML
ALL
CML
DIMT1 relative expression
8
6
4
2
0
B
GFP+
DIMT1
sgRNA
negative
selection
Cas9+ cancer
cell line
Day 3
Day 12
C
MOLM-13C
MOLM-13C DIMT WT
MOLM-13C DIMT E85A
MOLM-13C empty vector
Orange Fluorescence
Green Fluorescence
R2

**Fig. S1. Establishing DIMT1-expressing MOLM-13C cell lines. A.** Left, CERES dependency scores of DIMT1 in all cancers and different subtypes of leukemia. The CERES score is a computationally estimated gene dependency score based on CRISPR-Cas9 essentiality screens in different cancer cells. CERES accounts for the copy number specific effect and variable sgRNA activity. The lower the CERES score in the cell lines indicate a higher dependency on the gene of interest. Right, DIMT1 expression is slightly upregulated in leukemia cells. **B.** Schematic of CRISPR-based DIMT1 knockout in MOLM-13C cells. **C.** Flow cytometry of MOLM-13C cells re-expressing WT-DIMT1, E85A-DIMT1, or an empty vector before FACS sorting. Cells with Orange (mCherry)-fluorescence signals higher than $1 \times 10^2$ were collected.

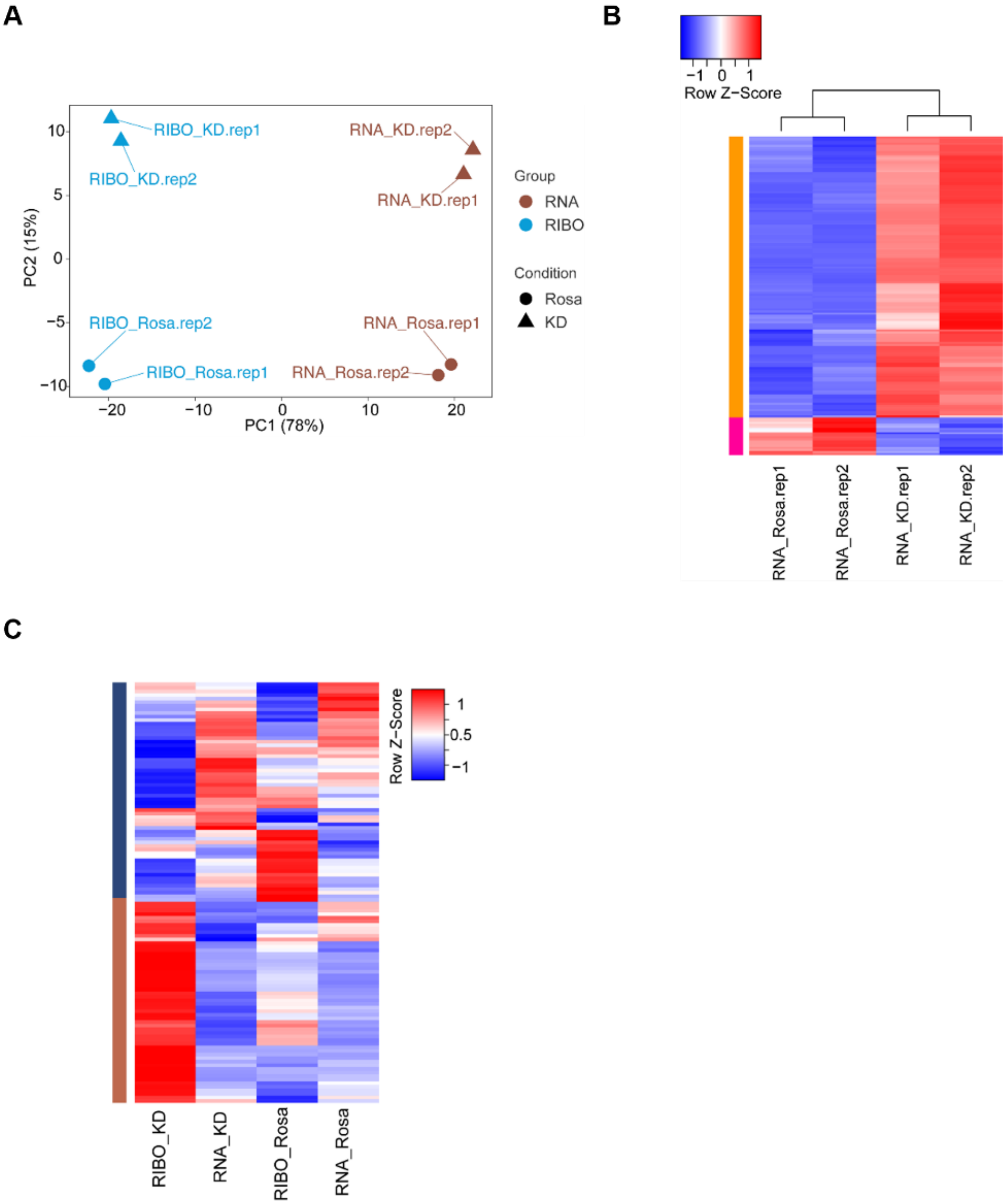


**Fig. S2. DIMT1 depletion alters the expression and translation efficiency of mRNAs. A.** PCA analysis of the ribo-seq samples (two biological replicates for each group and condition). The sequencing data of ribosome protected RNA (RIBO) and the sequencing data of the RNA input (RNA) from DIMT1 depletion, and the control cells separated well by sequencing group (RNA *vs.* RIBO) and treatment condition (Rosa *vs.* KD), and the two biological conditions cluster together, indicating the reliable quality of our sequencing data. **B.** Heatmap representing the read counts of the differentially expressed transcripts (as shown in Fig. 2a) of the RNA sequencing data. **C.** Heatmap representing the read counts of the differentially translated transcripts (as shown in Fig. 2d) of the RIBO and RNA sequencing data.

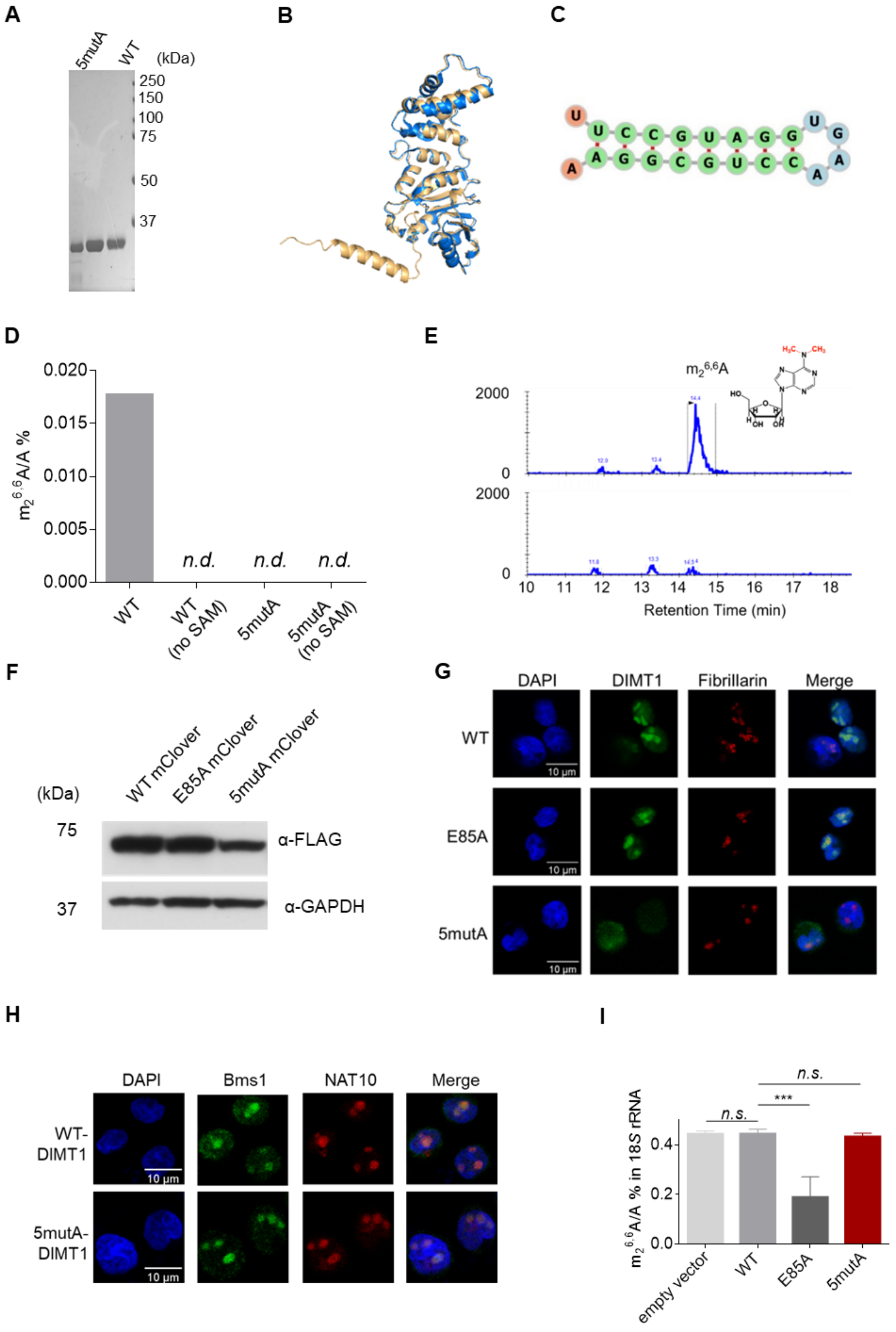
A
5mutA
WT
(kDa)
250
150
100
75
50
37
B
C
U
U C C G U A G G U G
A G G C G U C C A A
A
D
m$_2^{6,6}$A/A %
0.020
0.015
0.010
0.005
0.000
n.d.
n.d.
n.d.
WT
WT (no SAM)
5mutA
5mutA (no SAM)
E
m$_2^{6,6}$A
2000
0
2000
0
10
11
12
13
14
15
16
17
18
Retention Time (min)
F
(kDa)
WT mClover
E85A mClover
5mutA mClover
75
α-FLAG
37
α-GAPDH
G
DAPI
DIMT1
Fibrillarin
Merge
WT
E85A
5mutA
10 μm
H
DAPI
Bms1
NAT10
Merge
WT-DIMT1
5mutA-DIMT1
10 μm
I
m$_2^{6,6}$A/A % in 18S rRNA
0.0
0.2
0.4
n.s.
***
n.s.
empty vector
WT
E85A
5mutA

**Fig. S3. 5mutA-DIMT1 presents weaker RNA-binding affinity. A.** SDS-PAGE of purified WT-, E85A-, and 5mutA-DIMT1. **B**. Superimposition of AlphaFold-predicted structures of 5mutA-DIMT1 (yellow) and WT-DIMT1 (blue). **C.** Structural analysis of the RNA probe (which was used in Fig. 2C) was performed with RNA fold web server. **D**. LC-MS/MS quantification of $m_2^{6,6}A/A$ levels of the RNA probe in the in vitro methylation assay catalyzed by DIMT1. Error bars represent mean ± s.d., n = 6. n.d. means not detected. The ratio of enzyme to RNA was 1: 300, in reactions at 16°C for an overnight. **E**. LC-MS/MS channels and peak areas of guanosine and $m_2^{6,6}A$ for the *in vitro* methylation reaction. **F**. Western blots showing Flag-tagged WT-, E85A-, and 5mutA- DIMT1-mClover expression in *DIMT1*$^{+/-}$ HEK 293T cells. **G**. Representative fluorescence microscopy images of WT-, E85A-, and 5mutA-DIMT1-mClover in MOLM13C cells. Fibrillarin was stained as a nucleolar marker using an anti-fibrillarin antibody (red), while the nuclei were stained with a NuclearMask dye (blue). Scale bar, 10 µm. **H**. Representative fluorescence microscopy images of MOLM-13C expressing WT- or and 5mutA-DIMT1 after endogenous DIMT1 depletion using sgDIMT1 (post-transduction day 5). DIMT1-binding protein BMS1 (green) remains in the nucleolus in both WT- and 5mutA-DIMT1 cell lines after depletion of the endogenous DIMT1. NAT10 was used as a nucleolus marker (red), while the nuclei were stained with a NuclearMask dye (blue). Scale bar, 10 µm. **I**. LC-MS/MS quantification of $m_2^{6,6}A/A$ levels in 18*S* rRNA extracted from *DIMT1*$^{+/-}$ + empty vector, *DIMT1*$^{+/-}$ + WT-, *DIMT1*$^{+/-}$ + E85A-, *DIMT1*$^{+/-}$ + 5mutA-DIMT1 HEK 293T cells. Error bars represent mean ± s.d., n = 6. *n.s.* means not detected.

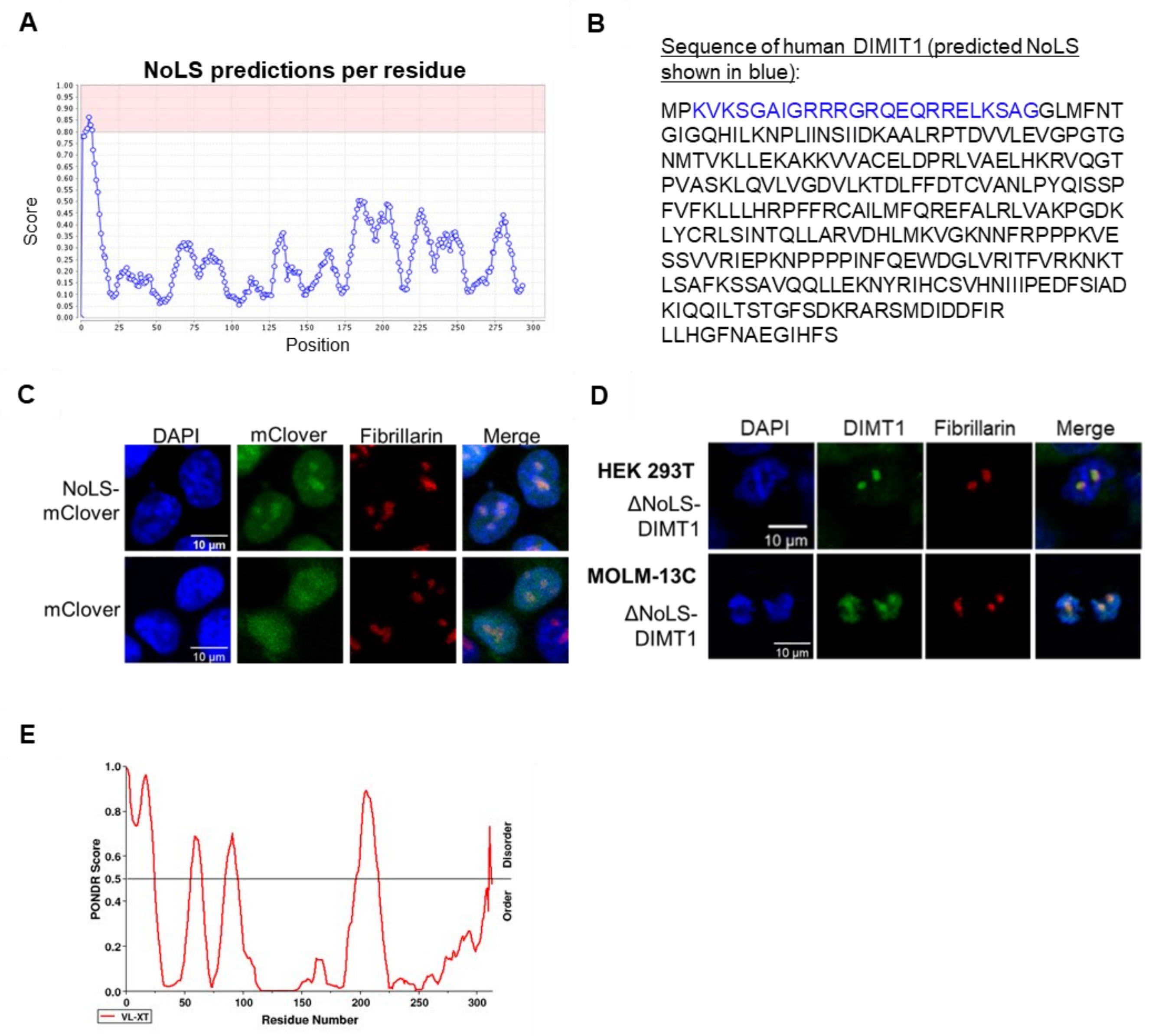


**Fig. S4. Prediction of the nucleolar-leading sequence (NoLS) in DIMT1. A.** NoLS prediction of DIMT1 was conducted using NoD software (http://www.compbio.dundee.ac.uk/www-nod/index.jsp). NoLSs are predicted regions of the protein in which at least 8 consecutive windows have an average score of at least 0.8. The NoLS score per residue graph identifies a segment of 20 residues representing 8 consecutive 13-residue windows (as explained in the predictor section). **B**. Amino-acid sequence of WT-DIMT1. The predicted NoLS is shown in blue.

**C.** Representative fluorescence microscopy images showing the localization of NoLS-mClover or mClover alone in HEK 293T cells. NoLS-mClover is a fusion protein with the predicted NoLS of DIMT1 fused to the N-terminus of mClover. The nucleoli were represented by fibrillarin (stained with an anti-fibrillarin antibody, red), while the nuclei were stained with a NuclearMask dye (blue). Scale bar, 10 μm. **D**. Representative fluorescence microscopy images of *DIMT1*$^{+/-}$ HEK 293T cells or MOLM-13C cells expressing ΔNoLS-DIMT1-mClover. The nucleoli were represented by fibrillarin (stained with an anti-fibrillarin antibody, red), while the nuclei were stained with a NuclearMask dye (blue). Scale bar, 10 μm. **E.** Disordered region prediction of DIMT1 amino acid sequence.

**Figure S5**

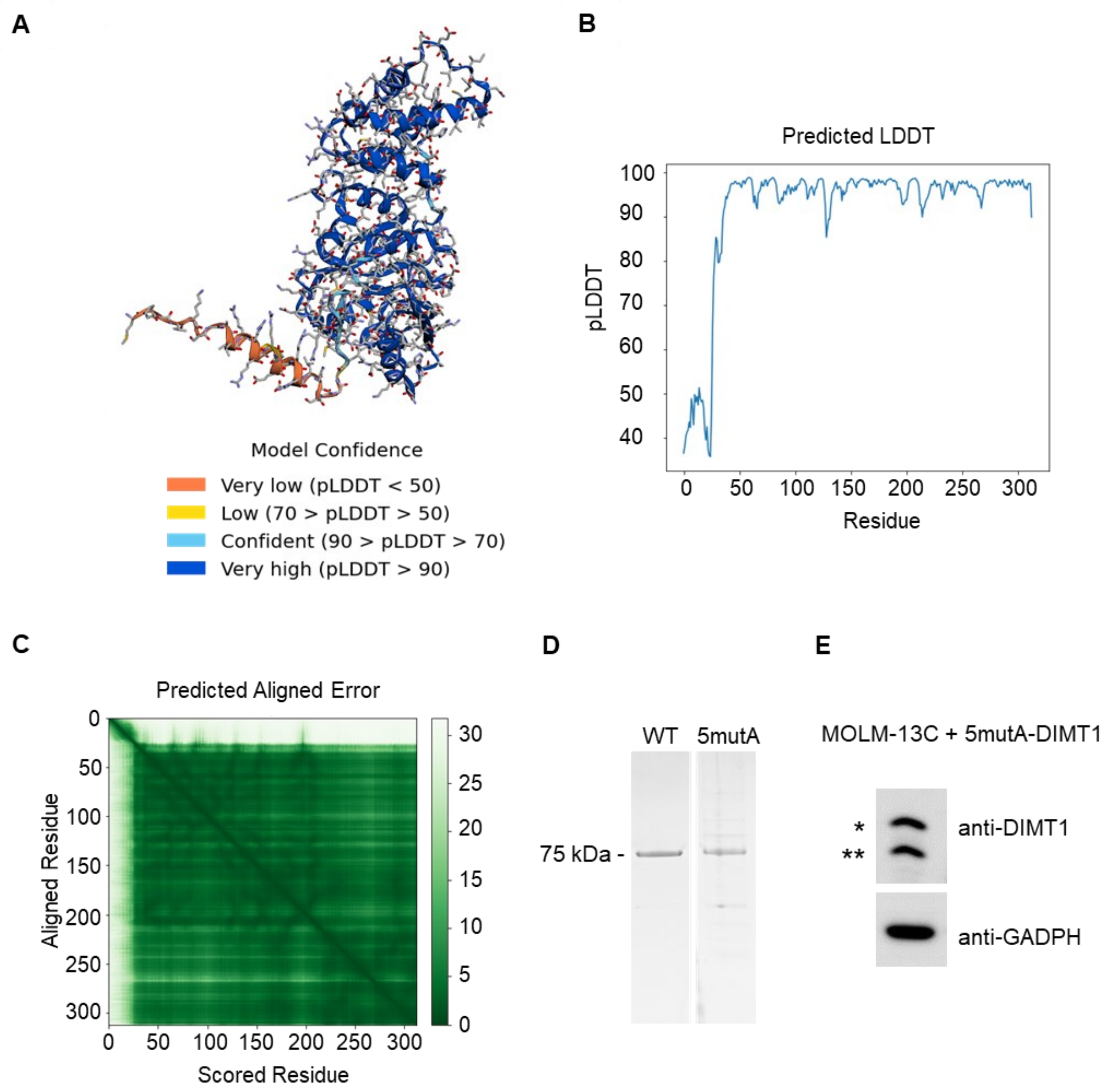


**Fig. S5. AlphaFold prediction of the structures of 5mutA-DIMT1. A.** AlphaFold predicted model of 5mutA with colors representing predicted local distance difference test (pLDDT) confidence. pLDDT determined based on standards of the inclusion radius determined by organizers of the critical assessment of techniques for protein structure prediction (CASP) biannual competition. **B.** Graph of pLLDT confidence across the amino acid sequence of 5mutA. **C.** Graph of predicted aligned error between every pair of amino acids within the sequence of

5mutA. **D**. SDS-PAGE of purified WT-DIMT1-mClover and 5mutA-DIMT1-mClover. **E.** Western blot showing endogenous DIMT1 (annotated by two stars) and exogenous DIMT1 (annotated by one star) expression in MOLM-13C cells expressing 5mutA-DIMT1.

**Table S1. High throughput sequencing data of the RNA-seq**

**Table S2. High throughput sequencing data of the ribo-seq**

**Table S3. Summary of the sequences of primers**
